# IFN-γ induces epithelial-to-mesenchymal transition of cancer cells via an unique microRNA processing

**DOI:** 10.1101/355263

**Authors:** U-Ging Lo, Rey-Chen Pong, Diane Yang, Leah Gandee, Elizabeth Hernandez, Andrew Dang, Chung-Jung Lin, John Santoyo, Shi-Hong Ma, Rajni Sonavane, Jun Huang, Shu-Fen Tseng, Loredana Moro, Arnaldo A. Arbini, Payal Kapur, Ganesh Raj, Dalin He, Chih-ho Lai, Ho Lin, Jer-Tsong Hsieh

**Author notes:** **Footnotes**: Jer-Tsong Hsieh, Ph.D.; 5323 Harry Hines Blvd., J8-134 Dallas, TX 75390, USA.

## Abstract

Interferon-γ (IFNγ) is a potent cytokine in modulating tumor immunity and tumoricidal effects. We demonstrate a new function of IFNγ in inducing epithelial-to-mesenchymal transition (EMT) in normal and cancer cells from different cell types. IFNγ activates JAK-STAT signaling pathway leading to the transcription of IFN-stimulated genes (ISGs), such as interferon-induced tetratricopeptide repeat 5 (IFIT5). We unveil a new function of IFIT5 complex in degrading precursor microRNAs (pre-miRNA) that include pre-miR-363 from the miR-106a-363 cluster, as well as pre-miR-101 and pre-miR-128 with a similar 5’-end structure with pre-miR-363. Noticeably, these suppressive miRNAs have similar functions by targeting EMT transcription factors in prostate cancer (PCa) cells. We further demonstrated that IFIT5 plays a critical role in IFNγ-induced cell invasiveness *in vitro* and lung metastasis *in vivo*. Clinically, IFIT5 is highly elevated in high-grade PCa and its expression inversely correlates with these suppressive miRNAs. Altogether, this study unveils pro-tumorigenic role of the IFN pathway via a new mechanism of action, which certainly raises concern about its clinical application.

## INTRODUCTION

Interferon-γ (IFNγ) is first characterized as a cytokine associated with antivirus as well as antitumor activities during cell-mediated innate immune response^1,2^. Mechanistically, IFNs can activate JAK-STAT signaling pathway after binding to type II receptors and induce the transcriptional activation of a variety of IFN-stimulated genes (ISGs) resulting in diverse biologic responses^3^. Among ISGs, interferon-induced tetratricopeptide repeat (IFIT) family members are highly inducible. They are viral RNA binding proteins^4^ and a part of antiviral defense mechanisms. They disrupt viral replication and/or viral RNA translation in host cells. Among IFIT orthologs, human IFIT1, IFIT2 and IFIT3 form a complex through the tetratricopeptide repeats (TPR) to degrade viral RNA. However, the functional role of IFIT5 is not fully understood since it acts solely as a monomer that can not only bind directly to viral RNA molecules via its convoluted RNA-binding cleft, but also endogenous cellular RNAs with a 5’-end phosphate cap, including transfer RNAs (tRNA) ^5,6^. In this study, we demonstrate a new function of IFIT5 in regulating microRNAs (miRNA) turnover.

miRNAs are a large family of short sequence single-stranded noncoding RNAs, which have been shown to regulate approximately 60% of protein-coding genes via post-transcriptional suppression, mRNA degradation, or translation inhibition^7,8^. Many miRNAs are associated with different stages of tumor development. These miRNAs are divided into onco-miRNAs and tumor suppressor miRNAs based on the function of their target genes. Similar to most protein-coding genes, miRNA genes can be regulated at transcriptional or post-transcriptional level ^9^. Unlike most eukaryotic protein genes, several miRNAs such as miR-106a-363^10^ and miR-17-92 are clustered together to generate a polycistronic primary transcript ^11–13^, which further complicates the regulatory scheme of miRNA biogenesis. For example, miR-363 belongs to the polycistronic miR-106a-363 cluster containing six miRNAs (miR-106a, miR-18b, miR-20b, miR-19b-2, miR-92a-2 and miR-363). Unlike the other five miRNAs with similar seed sequences and functions as the oncogenic miR-17-92 cluster ^14^, miR-363 acts as a tumor suppressor that is able to inhibit the transcriptional factor responsible for epithelial-to-mesenchymal transition (EMT). We further delineate that the differential regulation of miR-363 from this miR-106a-363 cluster is mediated by unique miRNA turnover machinery composed of IFIT5 and XRN1, which appears to be a novel, and previously unreported function of IFIT5.

Based on the discovery of IFIT5-elicied miR-363 degradation, additional miRNAs such as miR-101 and miR-128 are subjected to IFIT5 complex and these miRNAs are able to target several EMT transcriptional factors such as ZEB1 and Slug. Clinically, loss of these miRNAs is associated with tumor grade of PCa, which is inversely correlated with elevated IFIT5 mRNA level. On the other hand, IFIT5 mRNA expression is correlated with ZEB1 and Slug mRNA expression in PCa specimens. Functionally, IFIT5 plays a key role in IFNγ–induced EMT. Taken together, we conclude that IFN is a promoting factor in PCa progression.

## RESULTS

### The specific regulation of miR-363 expression

IFNγ is known to modulate cancer immunity and increase cytotoxicity. In this study, we observed that IFNγ was able to induce the expression of Slug, a potent EMT transcription factor, in PCa cell (C42) treated with in a dose-dependent manner (Fig. 1A). In contrast, reduced Disabled homolog 2-interacting protein (DAB2IP) protein expression was detected in treated cells (Fig. 1A). We have previously identified DAB2IP as an EMT inhibitor in PCa (PCa) ^15,16^. Thus, we believe that the mechanism of IFNγ-induced EMT is mediated through DAB2IP-regulated pathway. However, the inhibitory mechanism of DAB2IP in EMT is not fully characterized. Since emerging evidence demonstrates the critical role of miRNA in EMT process, we focused on the effect of DAB2IP on miRNA regulation. From miRNA microarray screening (Fig. S1A and B) between DAB2IP-proficient and–deficient cells, miR-363 is significantly decreased in DAB2IP-knockdown (KD) cells. The down-regulation of miR-363 in DAB2IP-KD cells was further validated in not only immortalized normal prostatic epithelial cell (RWPE1 and PNT1A) but also PCa lines (LAPC-4 and PC3) (Fig. 1B) and other cancer type such as renal cells (786O-KD and HK2-KD) (Fig. S1C). Ectopic expression of DAB2IP in C4-2Neo or LAPC4-KD cells (Fig. 1C), or HEK293 (Fig. S1D) was able to induce mature miR-363 levels in a dose-dependent manner, indicating that DAB2IP could modulate miR-363 expression.

**Figure 1.**
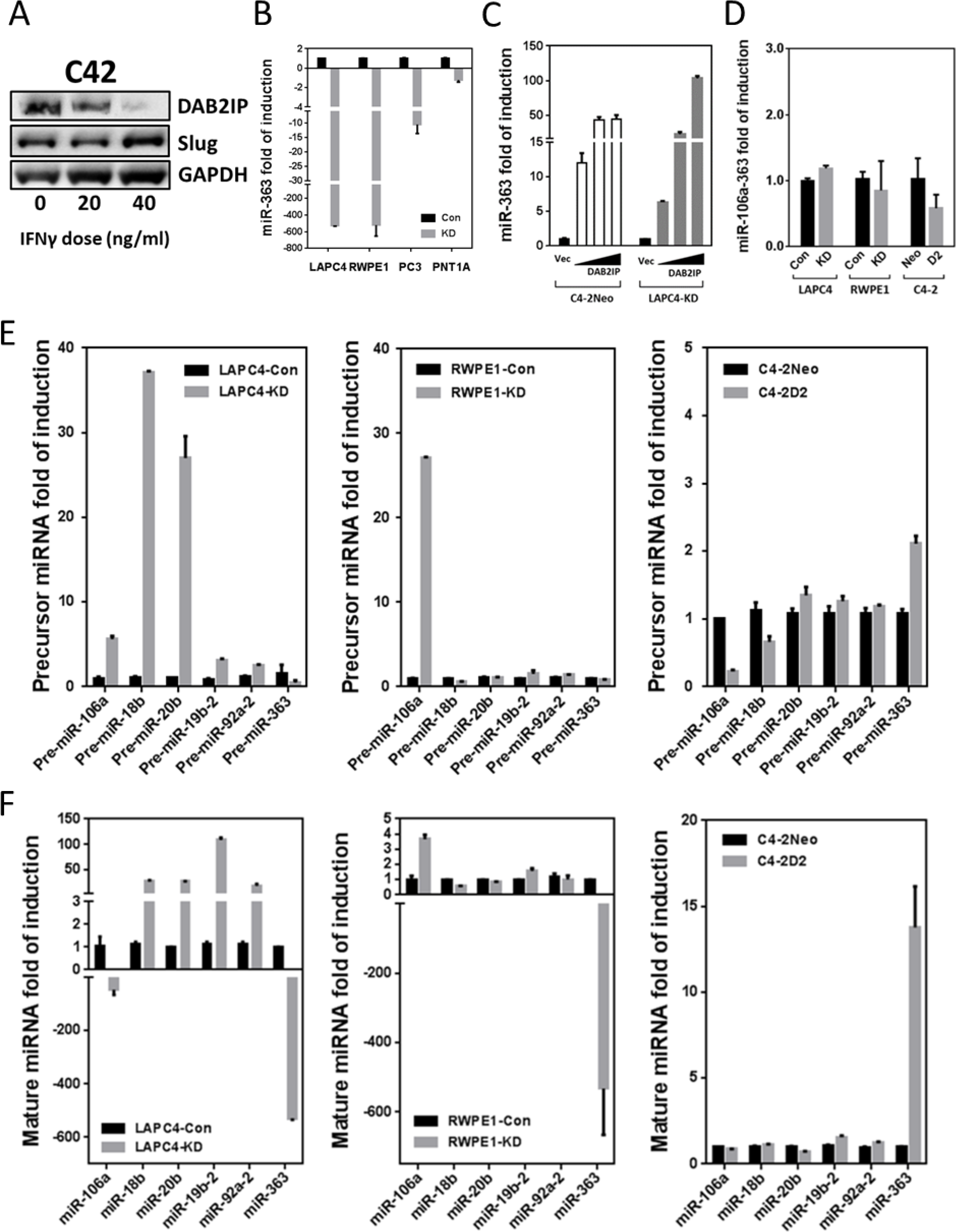
The effect DAB2IP on miR-363 expression in prostate cell lines. (A) Induced expression of DAB2IP and Slug protein level in C42 cells after treated with IFNγ for 24hrs. (B) Expression levels of miR-363 in DAB2IP-knockdown (KD) prostate cell lines after normalizing with the control (Con). (C) Induction of miR-363 by ectopic expression of DAB2IP in C4-2Neo and LAPC4-KD cell lines after normalizing with the control vector (Vec). (D) Expression levels of primary miR-106a-363 in DAB2IP-positive and-negative sublines. (E) Expression levels of precursor miRNAs (miR-106a, miR-18b, miR-20b, miR-19b-2, miR-92a-2 and miR-363) in DAB2IP-positive and-negative cells. (F) Expression levels of mature miRNAs (miR-106a, miR-18b, miR-20b, miR-19b-2, miR-92a-2 and miR-363) in DAB2IP-positive and-negative cells.

miR-363 is located in the polycistronic miR-106a-363 cluster that is first transcribed into a single primary miRNA containing the entire sequence of the cluster genes. We therefore examined the effect of DAB2IP on the expression levels of primary transcript of miR-106a-363. In contrast to the significant down-regulation of mature miR-363 in DAB2IP-KD cells, the expression levels of primary miR-106a-363 were similar between DAB2IP-positive and -negative cells (Fig. 1D). Also, the expression levels of pre-miR-363 showed no significant difference between these cells (Fig. 1E). Noticeably, only the mature miR-363 levels dramatically decreased in DAB2IP-KD cells (i.e., LAPC4-KD and RWPE1-KD) (Fig. 1F). On the other hand, only the mature miR-363 levels increased significantly in C4-2D (Fig. 1F) and HEK293D cells (Fig. S1E) with ectopic expression of DAB2IP. These findings indicate that DAB2IP specifically regulates miR-363 maturation process.

### The effect of IFIT5 on the distinct biogenesis of miR-363 from the miR-106a-363 cluster

In order to elucidate the machinery responsible for miR-363 maturation process, the protein candidates were immunoprecipitated using synthetic pre-miR-363 RNA molecule as bait and analyzed by LC-MS/MS (Table S1). This experiment unveiled IFIT5 protein as a potential hit. The steady-state levels of IFIT5 mRNA and protein were inversely correlated with DAB2IP (Fig. S2A). IFIT5 is characterized as a viral RNA or cellular tRNA binding protein and has, so far, not been known to bind to microRNAs. Therefore, the specific RNA-protein association between pre-miR-363 and IFIT5 was further validated using RNA pull-down and western blot analysis between DAB2IP-negative and–positive PCa cell lines (Fig. 2A). This inhibitory effect of DAB2IP on IFIT5 expression was further confirmed by the ectopic expression of DAB2IP in LAPC4-KD (Fig. 2B), C4-2Neo (Fig. 2C), LNCaP (Fig. S2B), and embryonic kidney cell (HEK293) (Fig. S2C).

To further examine whether IFIT5 plays a critical role in DAB2IP-mediated miR-363 maturation, we ectopically expressed IFIT5 in DAB2IP-positive cells. This resulted in a significant reduction of mature miR-363 levels but not the other mature miRNAs from the same cluster (Fig. 2D). In addition, on applying different IFIT5 small interfering RNA (siRNA) on LAPC4-KD cells, the levels of mature miR-363 correlated with the reduction in the endogenous IFIT5 mRNA levels (Fig. S2D). Thus, the IFIT5-C siRNA was chosen for further analysis. Reduced IFIT5 resulted in a significant elevation of mature miR-363 in a dose-dependent manner (Fig. S2E). Despite the significantly elevated mature miR-363 in IFIT5-KD cells, the levels of mature miR-106a, miR-18b, miR-20b, miR-19b-2 and miR-92a-2 remained relatively unchanged (Fig. 2E and S2F). Also, the expression levels of pre-miR-363 and other pre-miRNAs from the same cluster remained unchanged in these IFIT5-siRNA knockdown cells (Fig. S2G). Taken together, the data suggest that IFIT5 can specifically inhibit miR-363 maturation from the miR-106a-363 cluster.

**Figure 2.**
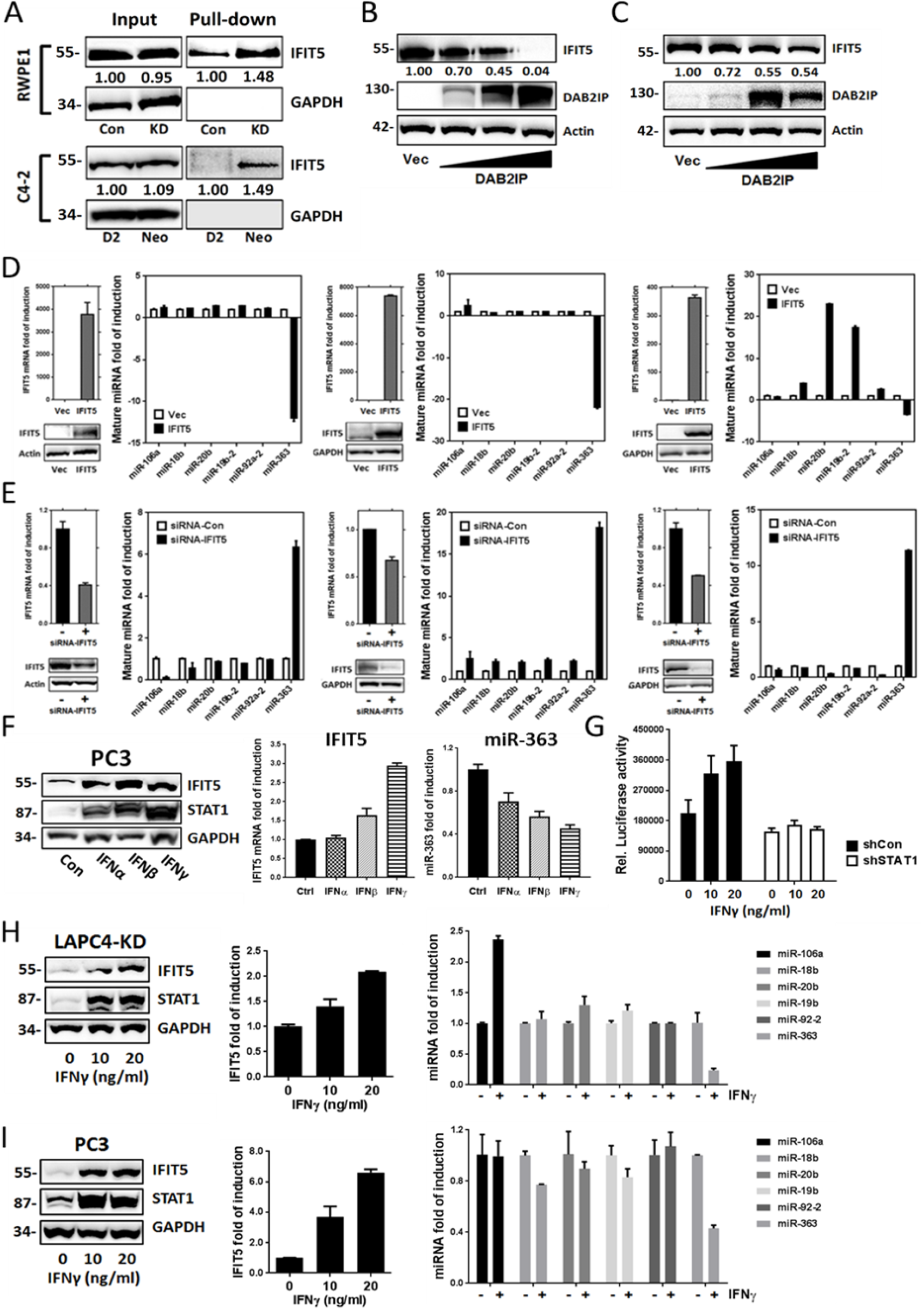
The impact of IFIT5 on miR-363 maturation from the miR-106a-363 cluster. (A) The interaction between IFIT5 protein and pre-miR-363 in DAB2IP-positive and – negative cells using RNA pull down assay. (B-C) Suppression of IFIT5 protein expression by ectopic transfecting DAB2IP into LAPC4-KD (B) and C4-2Neo (C) cells after normalizing with the control vector (Vec). (D) Expression levels of mature miRNAs (miR-106a, miR-18b, miR-20b, miR-19b-2, miR-92a-2 and miR-363) in IFIT5-overexpressing (IFIT5) LAPC4-Con, C4-2D2 and RWPE1-Con cells after normalizing with the control vector (Vec). (E) Expression levels of mature miRNAs (miR-106a, miR-18b, miR-20b, miR-19b-2, miR-92a-2 and miR-363) in IFIT5-siRNA knockdown (+) LAPC4-KD (left panel), C4-2Neo (middle panel) and RWPE1-KD (right panel) cells compared to the control siRNA (−). (F) Left and middle panel: Induction of IFIT5 protein and mRNA level by IFNα, IFNβ and IFNγ treatment for 48hrs in PC3 cells. Right panel: Expression level of miR-363 in PC3 cells treated with IFNα, IFNβ and IFNγ for 48hrs. (G) IFNγ-induced IFIT5 promoter activity in PC3 cells with shRNA knockdown of STAT1 (shSTAT1), compared to control shRNA (shCon). (H-I) Left and middle panel: Dose dependent induction of IFIT5 protein and mRNA level in LAPC4-KD and PC3 cells treated with IFNγ (10 and 20ng/ml) for 48 hrs. Right panel: Expression levels of mature miRNAs (miR-106a, miR-18b, miR-20b, miR-19b-2, miR-92a-2 and miR-363) in LAPC4-KD and PC3 cells treated with IFNγ (10 and 20 ng/ml) for 48 hrs.

### The effect of IFNs on the biogenesis of miR-363 from the miR-106a-363 cluster

Since the IFIT protein family is a typical ISG, we further confirmed that IFIT5 was induced in PC3 cells (PCa cell line) by all the IFNs (Fig. 2F). Meanwhile, a significant reduction of miR-363 was also observed under the same condition (Fig. 2F). Among three IFNs, IFNγ appears to have the most potent effect in inducing IFIT5 mRNA and suppressing miR-363. We therefore, used IFNγ to examine its impact on IFIT5 downstream target miRNAs. We first identified that the induction of IFIT5 mRNA by IFNγ was the result of transcriptional activation mediated by STAT1 signaling using IFIT5 gene promoter construct (Fig. 2G). Moreover, a dose-dependent induction of IFIT5 protein and mRNA by IFNγ was detected in other PCa cell line such as LAPC4-KD and PC3 (Fig. 2H and I) and significantly reduce mature miR-363 levels compared to other miRNAs in miR-106a-363 cluster (Fig. 2H and I, S2H and I). These data support a key mediator role of IFIT5 in IFNγ-mediated miR-363 suppression. Noticeably, the other member of IFIT family, such as IFIT1, was absent in PC3 cell after IFNγ treatment in contrast to RWPE1 cell (Fig. S2J), implying IFIT5 plays a unique role in PCa progression.

### The functional role of miR-363 in EMT

DAB2IP is known to inhibit EMT^16^. Based on the predicted sequences and gene profiling modulated by miR-363 in DAB2IP-KD cells, Slug/SNAI2 mRNA appears to be a potential candidate target gene. We therefore transfected miR-363-expression plasmid into KD cells, we observed huge expression levels (ranging 10,000 folds) then selected stable clone with low expression (<100 folds) to avoid any potential artifact. Indeed, suppression of Slug/SNAI2 mRNA levels was detected in miR-363 expressing cells compared to controls (Fig. 3A and B, and S3A). By constructing both wild type Slug/SNAI2 3’UTR (Slug-WT3’UTR) and mutant Slug/SNAI2 3’UTR (Slug-Mut^363^3’UTR) luciferase reporter genes, a significant reduction of the Slug-WT3’UTR but not the Slug-Mut^363^3’UTR activity was detected in RWPE-1-KD cells (Fig. 3C) and LAPC4-KD cells (Fig. S3B).

Slug/SNAI2 is known to promote EMT by suppressing the expression of epithelial markers such as E-Cadherin. As expected, an elevation of E-cadherin mRNA and protein was observed in miR-363 overexpressing RWPE-1-KD cells (Fig. 3D and E) and LAPC4-KD cells (Fig. S3C). In contrast, the expression of both mRNA and protein levels of Vimentin, a mesenchymal marker, were suppressed in both cell lines compared to controls (Fig. 3D and S3C). Functionally, miR-363 significantly reduced cell migration in miR-363 expressing RWPE-1-KD (Fig. 3F) and LAPC4-KD cells (Fig. S3D). In contrast, inhibition of miR-363 in RWPE-1 cells increased cell migration (Fig. 3G). We also noticed that cells collected from the lower chamber of a Transwell exhibited lower miR-363 levels than the upper chamber (Fig. S3D). Moreover, we restored Slug/SNAI2 level in miR-363-expressing cells that, in turn, resulted in a dose-dependent reduction of E-Cadherin mRNA and protein levels, as well as an elevation in Vimentin mRNA and protein levels in RWPE-1-KD cells (Fig. 3H) and LAPC4-KD cells (Fig. S3E). Moreover, we further observed an inverse relationship between miR-363 and IFIT5 expression in other cancer cell lines such as renal (such as 293T and 786O) and ovarian (such as HeLa) in addition to PCa cells (Fig. S4A). Knocking down of IFIT5 in these cell lines resulted in a significant elevation of mature miR-363 level (Fig. S4B). Similar inhibitory effect on EMT was observed in these cells: HeLa (Fig. S4C, D and E), 293T (Fig. S4F, G and H), and 786O cell lines (Fig. S4I, J, and K), by the ectopic expression of miR-363. These data indicate that miR-363 can suppress EMT by targeting the expression of Slug and Vimentin in cancer. Overall, examining the impact of miR-363 on EMT in different cancer cell lines further validates Slug/SNAI2 as the key target gene of miR-363-mediated EMT inhibition.

**Figure 3.**
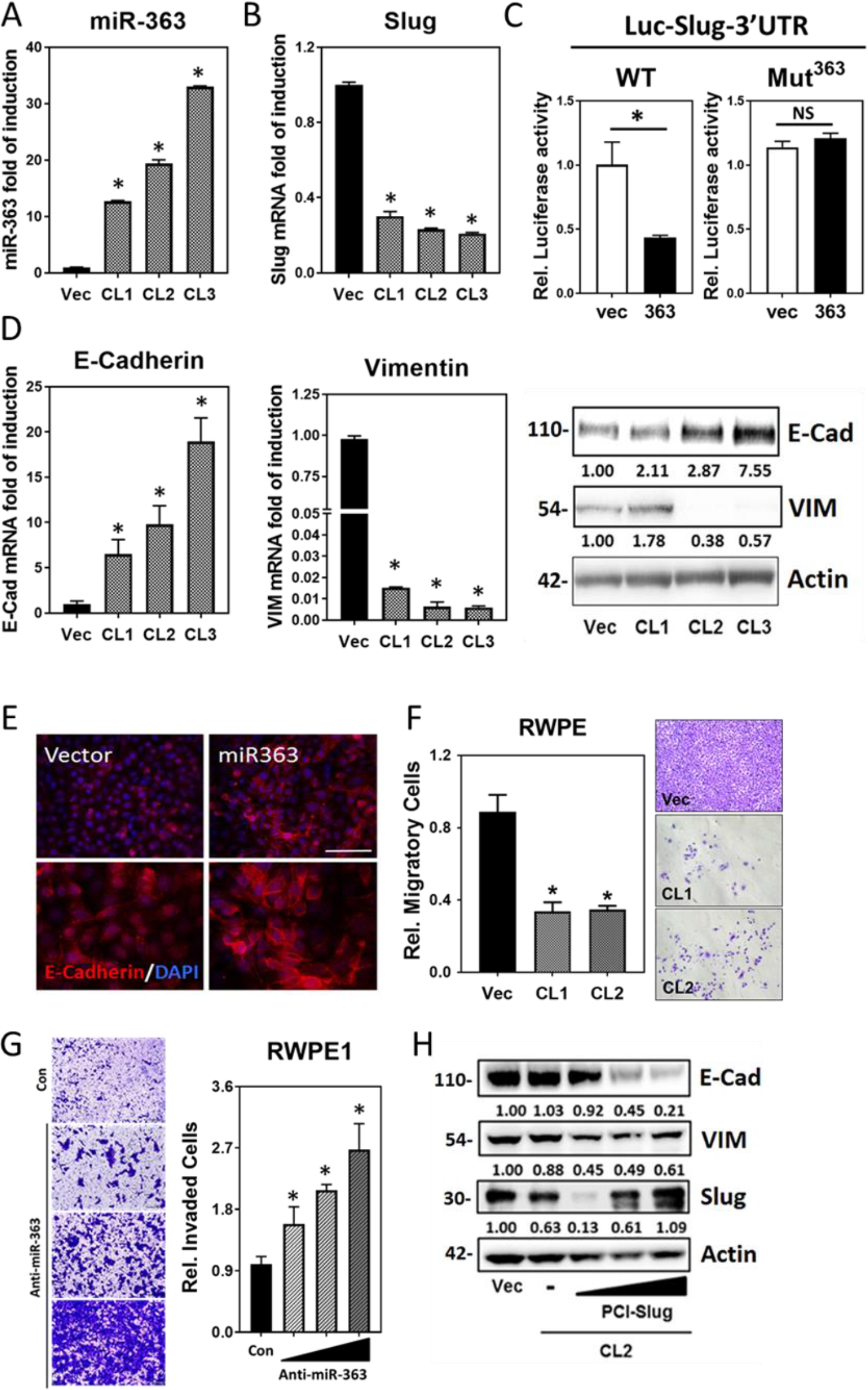
miR-363 reverse EMT signaling via targeting Slug. (A) Upregulation of miR-363 in RWPE1-KD cells transfected with pCMV-miR-363 plasmid. (B) Reduction of Slug mRNA expression and protein level in RWPE1-KD cells expressing miR-363. (CL: stable clone of RWPE1-KD cells expressing miR-363) (C) Luciferase reporter assay in RWPE1-KD cells co-transfected with siCHECK2-slug-WT 3’UTR or siCHECK2-Slug Mut363 3’UTR and pCMV-miR363 or empty vector. Luciferase activity unit is plotted as Renilla to Firefly luciferase activity (RFU). Each bar represents mean ± SD of four replicated experiments. (D) Induced mRNA and protein expression of E-cadherin and vimentin in RWPE1-KD cells expressing miR-363. (E) Immunofluorescence staining of E-cadherin protein expression in miR-363-overexpressed RWPE1-KD cells, compared to vector control (Scale bar = 100μm). (F) Transwell migration assay in RWPE1-KD cells-expressing miR363. Transmigrated RWPE1 cells were observed by crystal violet staining and quantified at OD 555nm. Each bar represents mean ± SD of three replicated experiments. (*<0.05). (G) Transwell invasion assay in RWPE1 cells transfected with anti-miR-363. Cells migrated to the lower bottom were stained with crystal violet and quantified at OD 555nm. Each bar represents mean ± SD of three replicated experiments. (*<0.05). (H) E-cadherin and Vimentin mRNA expression level after restoration of slug in RWPE1-KD cells-expressing miR363.

### The mechanism of IFIT5 on miR-363 turnover at precursor level

In a recent study, IFIT5 has been suggested to suppress virus replication by targeting the 5’-phosphate end of single stranded viral RNAs for rapid turnover^6^. Thus, we examined whether IFIT5 has a direct impact on the stability of pre-miR-363. In fact, pre-miR-363 RNA prepared from *in vitro* transcription was relatively stable at 37°C (Fig. S5A). However, the presence of IFIT5 protein complex significantly increased the turnover rate of pre-miR-363 RNA (Fig. S5A), indicating that the degradation of pre-miR-363 is accelerated by the IFIT5 protein complex. To examine the specificity of IFIT5 in the acceleration of pre-miR-363 degradation, we determined the *in vitro* degradation rate of pre-miR-92a-2 (immediate adjacent to miR-363) in the presence of IFIT5 and found no significant change (Fig. 4A). Previous studies ^4,5^indicate that IFIT5 protein binds to viral RNA molecules at either 5’-phosphate cap or 5’-tri-phosphate group. By comparing the 5’-end structure between pre-miR-92a-2 and pre-miR-363, we hypothesized that a single nucleotide (uracil) overhang in pre-miR-363, in contrast to the double-stranded blunt end in pre-miR-92a-2 is critical for IFIT5 recognition. Therefore, we generated 2 mutant pre-miR-363 constructs: one with 5’-end six nucleotides single stranded overhang (SS^6^Mut pre-miR-363) and the other with double-stranded blunt end (DSMut pre-miR-363) (Fig. 4B) to test their stabilities. The *in vivo* result (Fig. 4C) indicated that the expression levels of pre-miR363 or mature miR-363 derived from SS^6^Mut were significantly lower than those from native or DSMut form (Fig. 4C), indicating that the 5’-end structure of pre-miR-363 dictates the stability of miR-363 maturation. Similar pattern of mature miR-363 expression was also detected in RWPE1-KD (Fig. S4B). Upon determining the *in vitro* degradation rates of native, SS^6^Mut and DSMut pre-miR-363 RNA molecules, as we expected, SS^6^Mut pre-miR-363 was very sensitive to IFIT5 whereas DSMut pre-miR-363 was the most resistant one (Fig. 4D). Furthermore, we observed a steady elevation of SS^6^ Mut-derived mature miR-363 level in a dose-dependent manner in the presence of an incremental IFIT5 siRNA, while the expression of DSMut-derived mature miR-363 remained at high levels and was not affected by IFIT5 siRNA (Fig. 4E). Meanwhile, using RNA pull-down assay, SS^6^Mut pre-miR-363 exhibited higher affinity to IFIT5 protein than DSMut pre-miR-363 (Fig. 4F). Despite that these are artificial constructs, they are able to become mature *in vivo* form via miR biogenesis (Fig. 4C). As we expected, all these precursor constructs have very low binding affinity to Drosha (the ribonuclease to convert primary miR to precursor miR) (Fig. S5C). However, DSMut-pre-miR-363 exhibited higher binding to DICER (the ribonuclease to convert precursor miR to mature miR) than both native and SS^6^Mut pre-miR-363 (Fig. S4C), suggesting IFIT5 binding to native and SS^6^Mut pre-miR-363 could compete Dicer binding. Also, it appeared that DSMut pre-miR-363 was more stable than SS^6^Mut pre-miR-363 *in vivo* (Fig. 4C). Thus, comparing with native or SS^6^Mut pre-miR-363, DSMut exhibited a significant effect on inhibiting EMT (Fig. S5D) that is evidenced by elevated E-cadherin protein expression in RWPE1 cells (Fig. 4G). Also, DSMut exhibited a greater impact on diminishing cell migration of LAPC4-KD cells (Fig. S5E) and cell invasion of PC3 cells (Fig. 4H). These data conclude that IFIT5 recognizes the unique 5’-end overhanging structure of pre-miR-363 to elicit its degradation.

**Figure 4.**
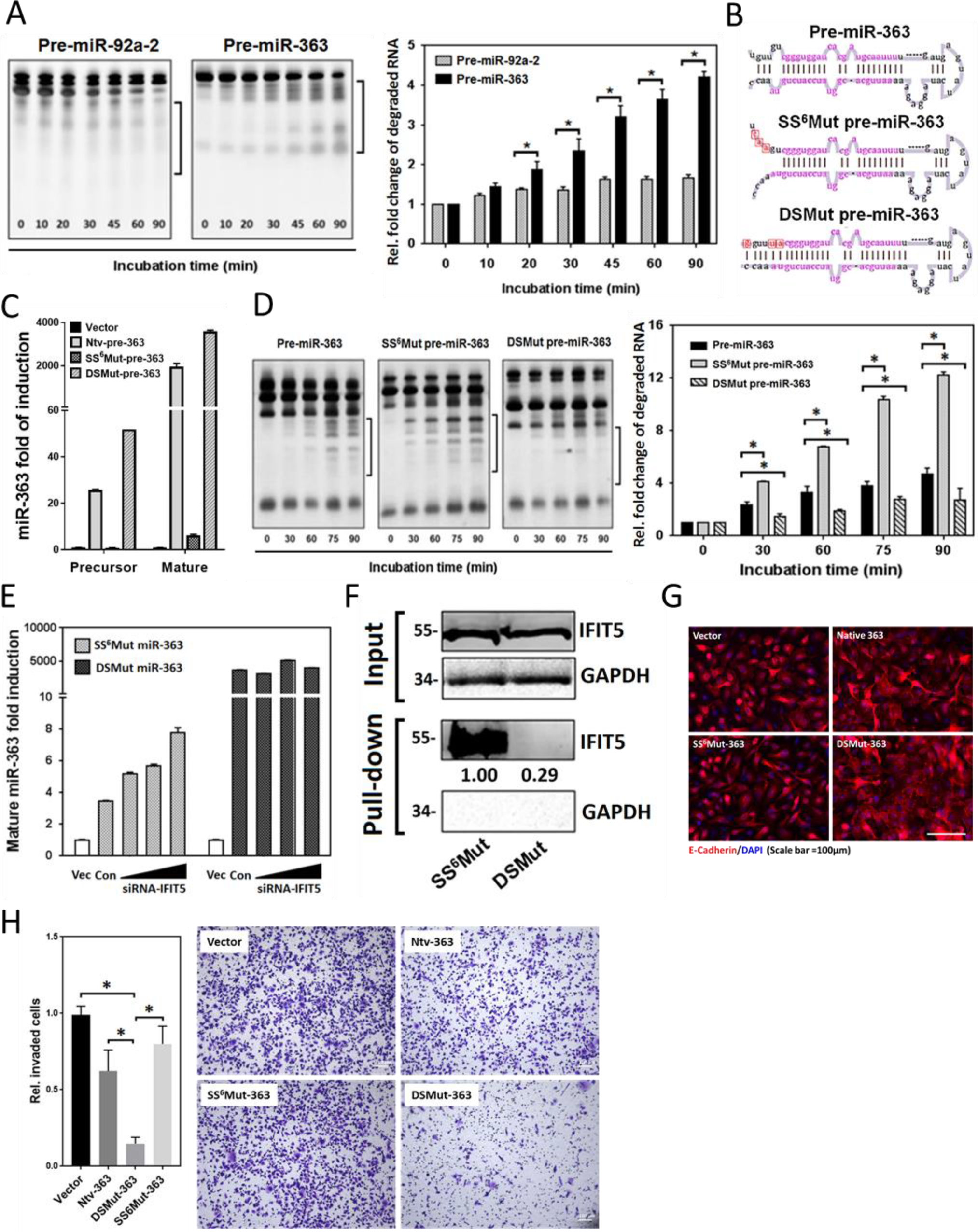
IFIT5-mediated precursor miR-363 degradation *in vitro.* (A) Time-dependent change of degraded pre-miR-92a-2 and pre-miR-363 fragments (bracket) after incubation with IFIT5 protein complex at 37°C normalized with 0 min. (*P<0.05) (B) Mutation of nucleotides (red box) for generating 5’-end 6 nucleotides single stranded pre-miR-363 (SS6Mut pre-miR-363) and blunt 5’-end double stranded pre-miR-363 (DSMut pre-miR-363). Both mature miR-363 and miR-363* sequence are shown in pink. (C) Expression levels of precursor and mature miR-363 in LAPC4-KD cells transfected with Native, SS6Mut or DSMut pre-miR-363 plasmids for 24 hrs after normalizing with the vector control. (D) Time-dependent change of degraded native, SS6Mut and DSMut pre-miR-363 fragments (bracket) after incubation with IFIT5 protein at 37°C, each time point was normalized with 0 min. (*p<0.05) (E) Induction of mature miR-363 in cells transfected with SS6Mut pre-miR-363 or DSMut pre-miR-363 plasmids and IFIT5 siRNA after normalizing with the control vector (Vec). (Con=control siRNA). (F) Interaction between IFIT5 protein and SS6Mut or DSMut pre-miR-363 RNA molecules using RNA pull down assay. (G) Immunofluorescence staining of E-cadherin protein expression in mutant pre-miR-363-overexpressed RWPE1 cells, compared to vector control. (H) The effect of Native, DSMut-or SS6Mut-pre-miR-363 on cell invasion in PC3 cells. Cells invaded at the lower bottom at the transwell were stained with crystal violet and counted. Each bar represents mean ± SD of nine fields of counted cell numbers. (* p<0.05)

To further demonstrate the specificity of this unique 5’-end structure of pre-miRNA, we also generated a mutant construct of pre-miR-92a-2 with single nucleotide at 5’-overhang (SS^1^Mut pre-miR-92a-2) (Fig. S5F) which is similar to the 5’-end of pre-miR-363 (Fig. 4B). Using RNA pull-down assay, we observed an increased interaction between SS^1^Mut pre-miR-92a-2 and IFIT5 protein, compared to native pre-miR-92a-2 (Fig. S5F). Moreover, the degradation rate of SS^1^Mut pre-miR-92a-2 increased in the presence of IFIT5 complex, compared to that of pre-miR-92a-2 (Fig. S5G). Thus, we believe that IFIT5-mediated precursor miRNAs turnover is determined by the 5’-end overhanging structure.

### The role of XRN1 in IFIT5-mediated miR-363 turnover

Although IFIT5 can elicit miR-363 turnover, IFIT5 doesn’t possess ribonuclease activity. To determine if a ribonuclease is associated with the IFIT5-pre-miR-363 complex, we further examined LC-MS/MS results and identified an exoribonuclease candidate-XRN1. XRN1 is known to regulate mRNA stability via cleavage of de-capped 5’-monophosphorylated mRNA ^17,18^ and a recent study also implied its potential role in miRNA turnover ^19^. Indeed, an interaction was observed between IFIT5 and XRN1 protein in LAPC4-Con cells transfected with Flag-tagged IFIT5 (Fig. 5A). Using three different XRN1 siRNAs in LAPC4-KD cells, we showed that the elevated expression levels of miR-363 correlated with the diminished level of XRN1 protein (Fig. 5B and S6A). Similar to IFIT5-knockdown, data from XRN1-KD cells clearly demonstrated that only mature miR-363 exhibited a significant accumulation whereas the levels of other mature miRNAs (miR-106a, miR-18b, miR-20b, miR-19b-2, and miR-92a-2) remained relatively unchanged (Fig. 5C).

**Figure 5.**
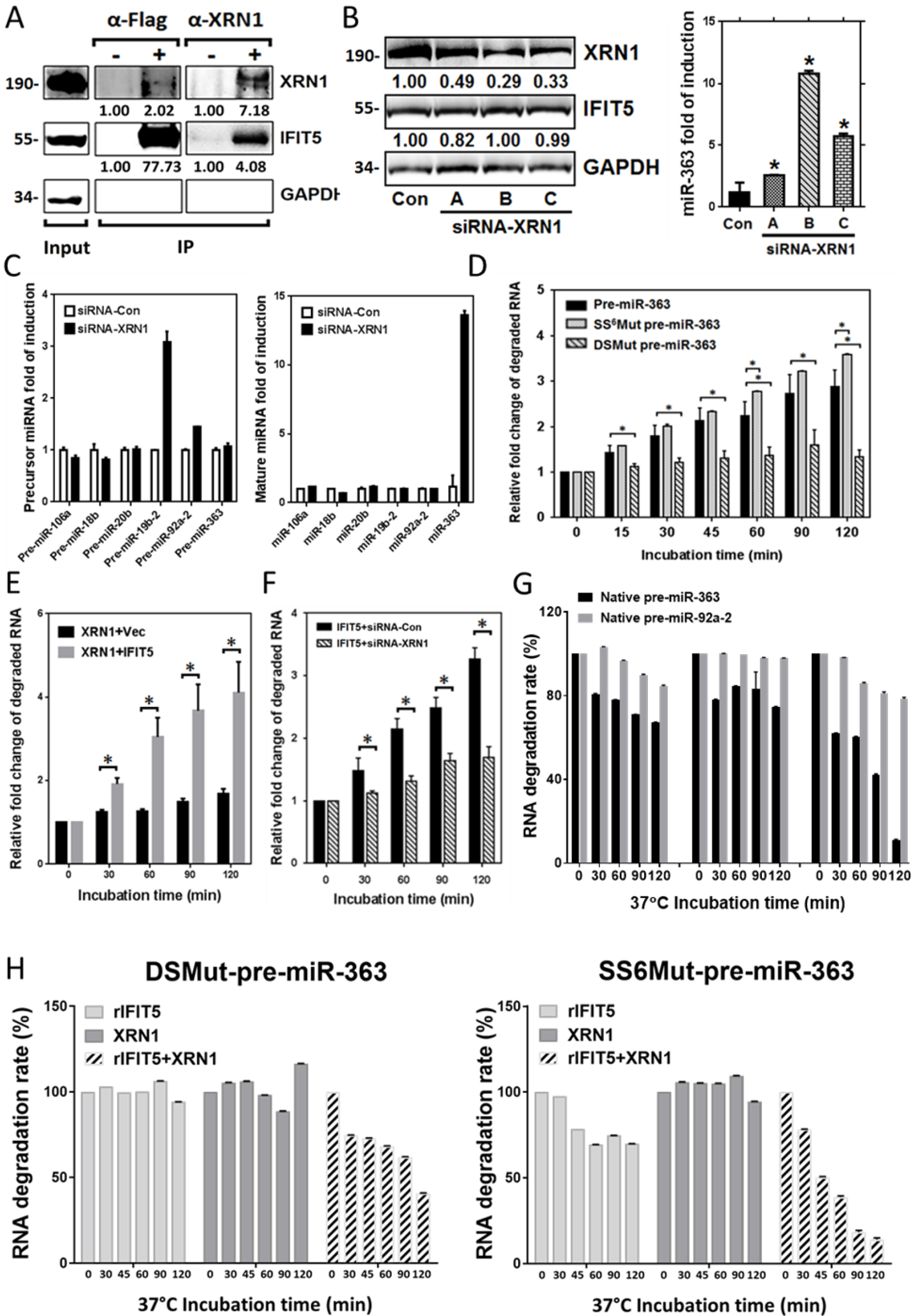
Interaction between XRN1 with IFIT5 leading to pre-miR-363 degradation *in vitro*. (A) Interaction between IFIT5 and XRN1 proteins using IP by Flag and XRN1 antibodies, respectively. (B) Left: knockdown of XRN1 in LAPC4-KD cells using siRNA. Right: Induction of mature miR-363 in LAPC4-KD cells transfected with XRN1 siRNA after normalizing with the control siRNA (Con). (C) Expression levels of precursor and mature miRNAs (miR-106a, miR-18b, miR-20b, miR- 19b-2, miR-92a-2 and miR-363) in XRN1-knockdown (siRNA-XRN1) LAPC4-KD cells after normalizing with the control siRNA (siRNA-Con). (D) Time-dependent change of degraded native, SS6Mut and DSMut pre-miR-363 fragments after incubation with immunoprecipitated XRN1 protein at 37°C after normalizing with 0 min. (*p<0.05) (E) Time-dependent change of degraded SS6Mut pre-miR-363 fragments after incubation with immunoprecipitated-XRN1 alone (XRN1+Vec) or XRN1-IFIT5 complex (XRN1+IFIT5) at 37°C after normalizing with 0 min. (*p<0.05) (F) Time-dependent change of degraded SS6Mut pre-miR-363 after incubation with the immunocomplex derived from cells transfected with IFIT5 and control siRNA (IFIT5+siRNA-Con) or XRN1 siRNA (IFIT5+siRNA-XRN1) at 37°C after normalizing with 0 min. (*p<0.05) (G) Degradation of native pre-miR-363 or pre-miR-92a-2 after incubation with recombinant IFIT5 protein (rIFIT5), XRN1 enzyme (XRN1) or combination of XRN1 and rIFIT5 at 37°C after normalizing with 0 min. (H) Degradation of SS6Mut-or DSMut-pre-miR-363 after incubation with rIFIT5, XRN1 or combination of XRN1 and rIFIT5 at 37°C after normalizing with 0 min.

Also, by incubating XRN1 immunocomplex (Fig. S6B) with native, SS^6^Mut or DSMut pre-miR-363 RNA *in vitro*, a significant increase of both native and SS^6^Mut pre-miR-363 degradation was detected in a time-dependent manner, whereas DSMut pre-miR-363 levels remained relatively unchanged (Fig. 5D and S6B), implying that the IFIT5 binding structure in the 5’-end of pre-miR-363 is critical for recruiting XRN1. In addition, by increasing IFIT5 expression in XRN1-expressing LAPC4-Con cells, XRN1-IFIT5 immunocpmplex apparently increased the *in vitro* degradation of SS^6^Mut-pre-miR-363 compared with control (XRN1 alone) (Fig. 5E and S6C). On the other hand, knocking down XRN1 in IFIT5-overexpressing LAPC4-Con cells diminished the *in vitro* degradation rate of SS^6^Mut pre-miR-363 after incubation with IFIT5 (Fig. 5F and S6D). These findings provide further evidence for the specific function of IFIT5-XRN1 complex in miR-363 turnover. In addition, using recombinant IFIT5 protein with or without XRN1 enzyme, the result (Fig. 5G and S6E) clearly indicated that both IFIT5 and XRN1 proteins are required to degrade pre-miR-363 transcript *in vitro*. Meanwhile, combination of recombinant IFIT5 and XRN1 shows less impact on the degradation of pre-miR-92a-2 (Fig. 5G and S6E). Similarly, the SS^6^Mut pre-miR-363 is more sensitive to rIFIT5-XRN1 complex-mediated degradation than DSMut pre-miR-363 (Fig. 5H and S6F). Overall, these data demonstrate that the IFIT5-XRN1 complex is responsible for the degradation of pre-miR-363.

### The effect of IFNγ on miR-101, miR-128, and miR-363 processing mediated by IFIT5

To survey additional miRNAs subjected to IFIT5-mediated precursor miRNA degradation, we performed miRNA microarray using IFIT5-expressing LAPC4-Con and IFIT5-depleted LAPC4-KD cells and unveiled miR-101 and miR-128 as candidates. We further confirmed that the presence of IFIT5 reduced the expression of mature miR-101 and miR-128 as well as miR-363 in PC3 cell line (Fig. 6A). In contrast, IFIT5-knockdown in LAPC4-KD cells increased the expression of all three miRNAs (Fig S7A). Also, XRN1 knockdown in IFIT5-expressing cells could rescue the expression levels of mature miR-363, miR-101 and miR-128 (Fig. S7B), indicating the requirement of XRN1 in IFIT5 complex in degrading these miRNAs. Similarly, IFNγ treatment resulted in reducing the expression of miR-101, miR-128 and miR-363 (Fig. 6B). This inhibitory effect of IFNγ can be reversed or diminished by knocking down IFIT5, STAT1 or XRN1 (Fig. 6C). Similarly, overexpression of DAB2IP in PC3 cells also diminished the inhibitory effect of IFNγ on the suppression of miR-101, miR-128 and miR-363 level (Fig. S7C), supporting the key role of IFIT5 in IFNγ-elicited precursor miRNAs processing.

**Figure 6.**
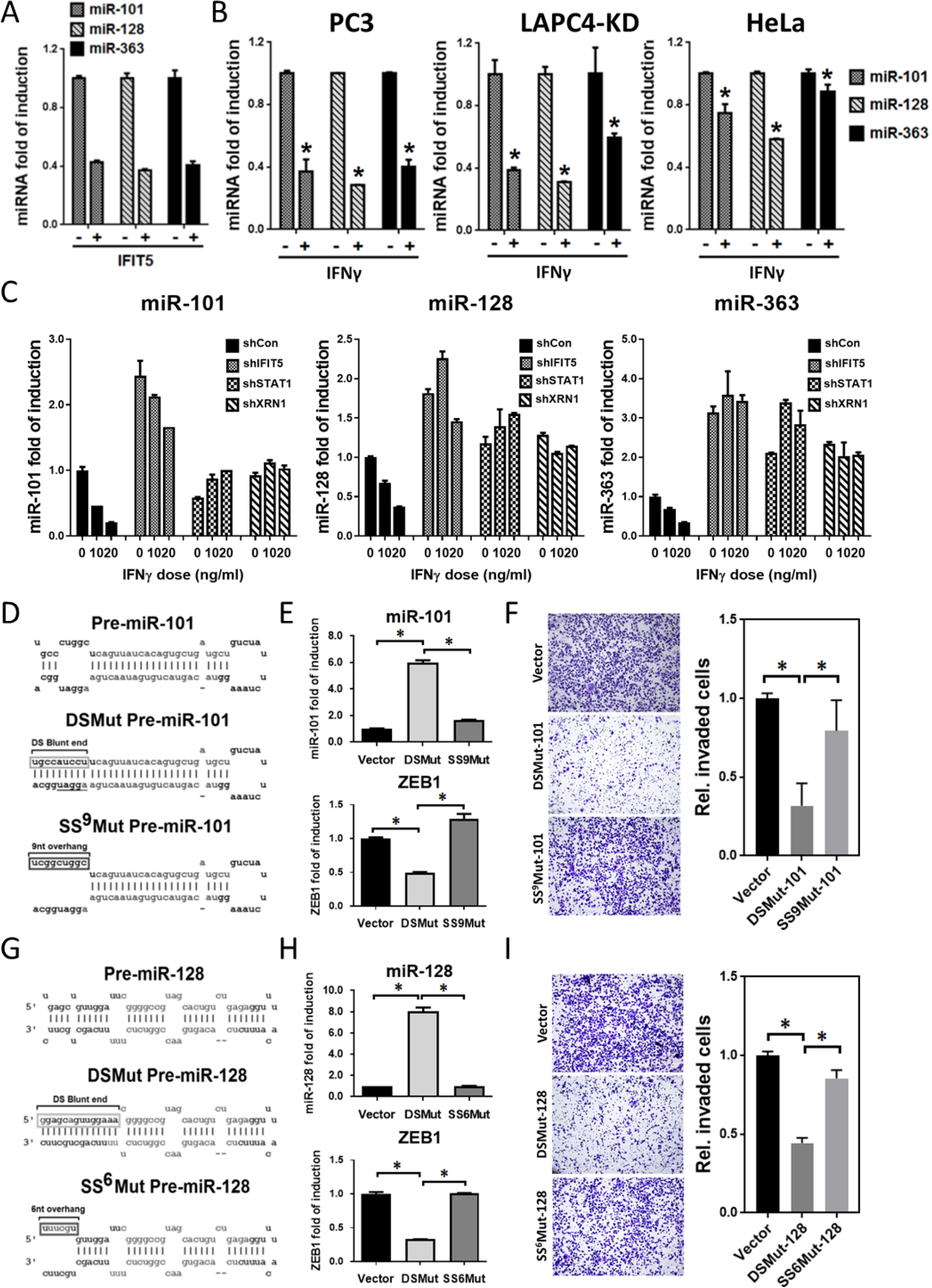
The impact of IFIT5-XRN1 mediated precursor miRNA degradation on EMT. (A) Expression level of mature miR-101, miR-128 and miR-363 in IFIT5-overexpressed PC3 cells. (B) Expression level of miR-101, miR-128 and miR-363 in PC3, LAPC4-KD and HeLa cells treated with IFNγ(+), compared with control vector (−). (C) Expression level of miR-101, miR-128 and miR-363 in PC3 cells treated with 0, 10 and 20 ng/ml dose of IFNγ after knockdown of IFIT5 (shIFIT5), STAT1 (shSTAT1) or XRN1 (shXRN1), compared to vector control (shCon). (D) Mutation of nucleotides (box) for generating blunt 5’-end double stranded pre-miR-101 (DSMut pre-miR-101) and 5’-end 9 nucleotides single stranded pre-miR-101 (SS9Mut pre- miR-101). Both mature miR-101 and miR-101* sequence are shown in lighter gray. (E) The effect of DSMut or SS9Mut pre-miR-101 on the expression level of mature miR-101 and ZEB1 mRNA (* p<0.05). (F) The effect of DSMut or SS9Mut pre-miR-101 on the cell invasion in PC3 cells. Cells invaded at the lower bottom at the transwell were stained with crystal violet and counted. Each bar represents mean ± SD of nine fields of counted cell numbers. (* p<0.05). (G) Mutation of nucleotides (box) for generating blunt 5’-end double stranded pre-miR-128 (DSMut pre-miR-128) and 5’-end 6 nucleotides single stranded pre-miR-128 (SS6Mut pre- miR-128). Both mature miR-128 and miR-128* sequence are shown in lighter gray. (H) The effect of DSMut or SS6Mut pre-miR-128 on the expression level of mature miR-128 and ZEB1 mRNA (* p<0.05). (I) The effect of DSMut or SS6Mut pre-miR-128 on the cell invasion in PC3 cells. Cells invaded at the lower bottom at the transwell were stained with crystal violet and counted. Each bar represents mean ± SD of nine fields of counted cell numbers. (* p<0.05).

By comparing the precursor structures of miR-101 and miR-128, it appeared that both pre-miR-101 and pre-miR-128 have similar 5’-end structure with pre-miR-363, we therefore generated 2 mutant constructs: one with 5’-end single stranded overhang (SSMut) and the other with double-stranded blunt end (DSMut) (Fig. 6D and G) to test their expression in IFIT5-expressing PC3 and LAPC4-KD cell lines. As we expected, DSMuts were resistant to IFIT5-elicited miRNA degradation and resulted in elevated expression of mature miRNA in PC3 (Fig. 6E and H) and LAPC4-KD cells (Fig. S7E and S7F). Based on the 3’UTR sequence, ZEB1 mRNA was predicted as a target for both miR-101 and miR-128 (Fig. S7D), and the results indeed indicated that both miR-101 and miR-128 could suppress ZEB1 mRNA levels (Fig. S7D). Again, DSMuts appeared to degrade ZEB1 mRNA and protein more efficiently in PC3 (Fig. 6E and H) and LAPC4-KD cells (Fig. S7E and S7F), which are correlated with the suppression of cell invasion in PC3 cells (Fig. 6F and I) and cell migration in LAPC4-KD cells (Fig. S7E and S7F). Based on the accumulating evidence that miR-200 family is one of the well-known miRNA groups in suppressing EMT factors such as ZEB1 and Slug, we also examined the effect of IFIT5 or IFNγ on the expression level of miR-200 family members involved in the suppression of Slug and ZEB1. The result (Fig. S7G) indicated that the impact of IFNγ or IFIT5 on EMT was not mediated through miR-200 family members. Overall, the effect of IFIT5-XRN1 complex on pre-miR-101/128/363 processing is unique with respect to the similar 5’-end overhang structure.

### The effect of IFNγ on EMT mediated by IFIT5

Based on the mechanism of action of IFIT5-XRN1 complex in the degradation of miRNAs that can target EMT factors, we further examined whether IFNγ could elicit EMT by suppressing these miRNAs via STAT1 signal axis and its downstream effector-IFIT5/XRN1complex. Indeed, IFNγ treatment increased the PC3 cell migration (Fig. 7A) and invasion (Fig. 7B) that was diminished in the absence of IFIT5 (Fig. 7A, 7B and S8A) or STAT1 (Fig. 7B, S8A and S8B), which is consistent with the expression of EMT factors (Slug and ZEB1) or decrease in the mesenchymal marker (Vimentin) or increase in the epithelial marker (E-cadherin) (Fig. 7C and D). As we expected the expression of all these three miRNAs was inhibited by IFNγ in a dose-dependent manner (Fig. 6C) and IFNγ failed to suppress the expression of these miRNAs in the absence of XRN1, STAT1 or IFIT5 (Fig. 6C) in which no induction of Slug and ZEB1 mRNA was detected (Fig. 7E). Similarly, the effect of IFNγ on Slug and ZEB1 mRNA induction can be diminished in cells transiently transfected with miR-101, miR-128 or miR-363 (Fig. S8C).

**Figure 7.**
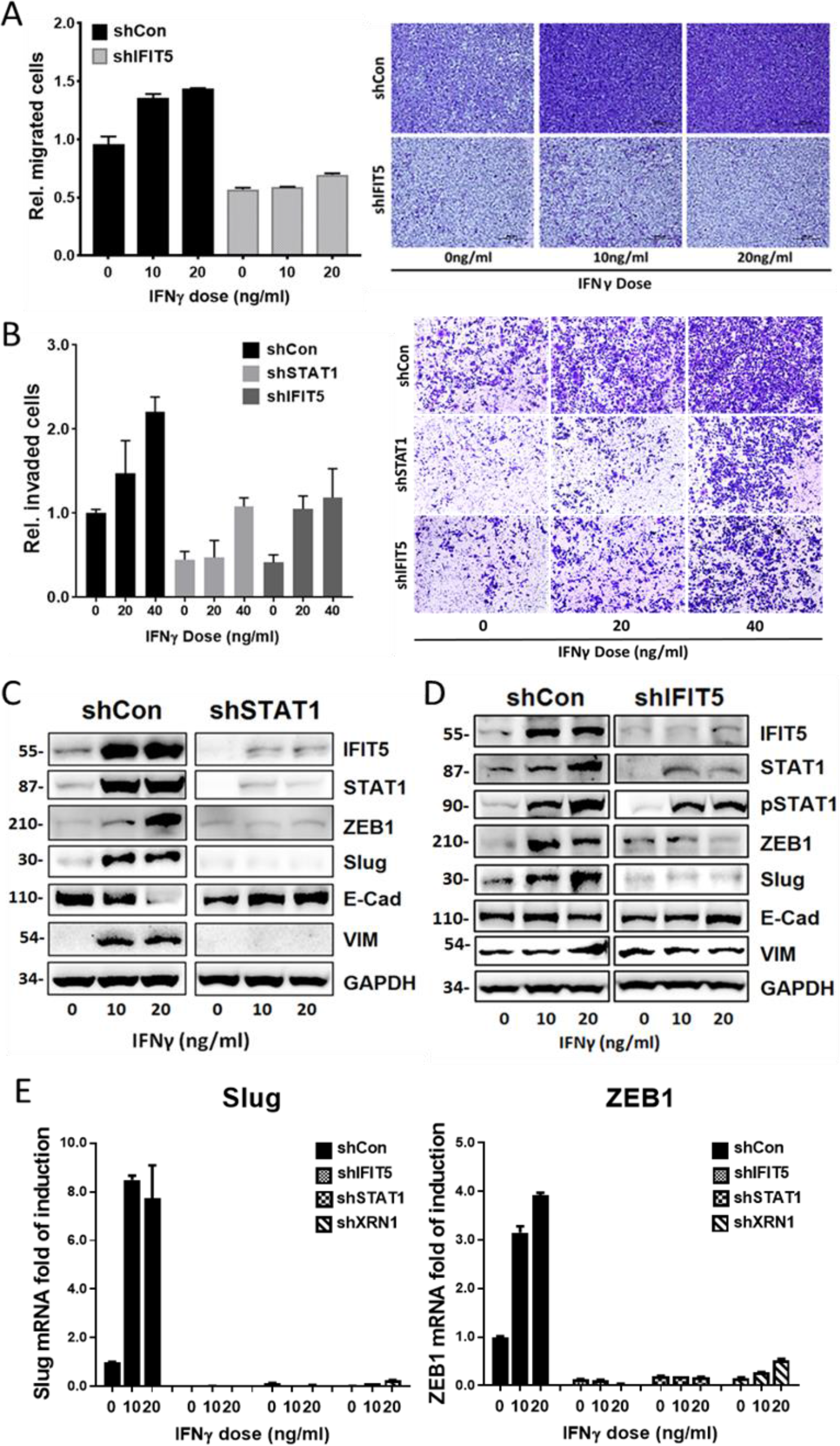
IFNγ elicits its impact on EMT via activating IFIT5-XRN1 mediated miRNA turnover through STAT1 signaling pathway. (A) Transwell migration of IFIT5-knockdown (shIFIT5) PC3 cells after treatment of IFNγ for 48hrs, compared to vector control (shCon). Migrated cells were stained with crystal violet and quantified at OD 555nm. Each bar represents mean ± SD of three replicated experiments. (* p<0.05, NS=no significance). (B) Transwell invasion of STAT1-or IFIT5-knockdown (shSTAT1, shIFIT5) PC3 cells after treatment of IFNγ (0, 20, and 40 ng/ml) for 48hrs, compared to control vector (shCon). Cells invaded at the lower bottom of transwell were stained with crystal violet and counted. Each bar represents mean ± SD of counted cell numbers from nine fields. (* p<0.05) (Scale bar=100μm). (C-D) Induction of IFIT5, E-cadherin and mesenchymal factors (ZEB1, Slug and Vimentin) in STAT1-or IFIT5-knockdown (shSTAT1 or shIFIT5) PC3 cell lines in response to IFNγ treatment, compared to PC3 cells with control vector (shCon). (E) Induction level of Slug and ZEB1 mRNA in PC3 cells treated with 0, 10 and 20 ng/ml dose of IFNγ after knockdown of IFIT5 (shIFIT5), STAT1 (shSTAT1) or XRN1 (shXRN1), compared to vector control (shCon).

Apparently, IFNγ is capable of inducing EMT at low concentrations that are not anti-tumorigenic or anti-proliferation (Fig. S8D); its direct anti-tumor activity is known at much higher concentration (>1000 ng/ml)^20^. These data provide new evidence that IFNγ is a potent inducer of EMT via STAT1-IFIT5/XRN1 signal axis of miRNA regulation.

### The clinical correlation of IFIT5, miRNAs and EMT biomarkers in PCa

To examine the *in vivo* effect of IFNγ on PCa metastasis and the role of IFIT5 in this event, we treated control (shCon) and IFIT5-knockdown cells with IFNγ for 48hrs then cells were injected intravenously into SCID animal via tail vein. IFNγ treatment significantly increases the number and size of metastatic nodules at lung parenchyma, in contrast, the loss of IFIT5 dramatically reduces metastasis of PCa with or without IFNγ (Fig. 8A, 8B and Table S2). Furthermore, we demonstrated the effect of IFNγ on EMT clinically, we employed an *ex vivo* culture system^21^ using human PCa specimens and data indicated that IFNγ was able to induce the expression of IFIT5, ZEB1, Slug (Fig. 8C) and Vimentin (Fig. S9A) gene whereas miR-101, miR-128 and miR-363 levels were significantly inhibited (Fig. S9B). We also surveyed the expression status of IFIT5 from different grades of PCa specimens and data (Fig. 8D) indicate that IFIT5 mRNA levels were significantly elevated in the high-grade PCa. As expected, the expression pattern of miR-363, miR-101 and miR-128 levels was opposite to that of IFIT5 (Fig. 8D), which is consistent with our observation from tissue culture cell lines. In contrast, miR-92a-2 and miR-19b-2 immediately adjacent to miR-363 known as oncomirs exhibited an elevated expression pattern in PCa tissues compared to normal tissues (Fig. S9C), supporting the specificity of IFIT5 on miRNA degradation. Meanwhile, data from PCa specimens also demonstrated a similar correlation between IFIT5 mRNA and miR-363 or miR-101 (Fig. 8E). In addition, analyses of expression of EMT factors or markers demonstrated a positive correlation between IFIT5 and ZEB1 (or Slug) (Fig. 8F), and Vimentin (Fig. S9D). Similar correlation between IFIT5 and ZEB1 (or Slug) was also observed in TCGA dataset of renal cell carcinoma (Fig. S9E) and ovarian carcinoma (Fig. S9F).

**Figure 8.**
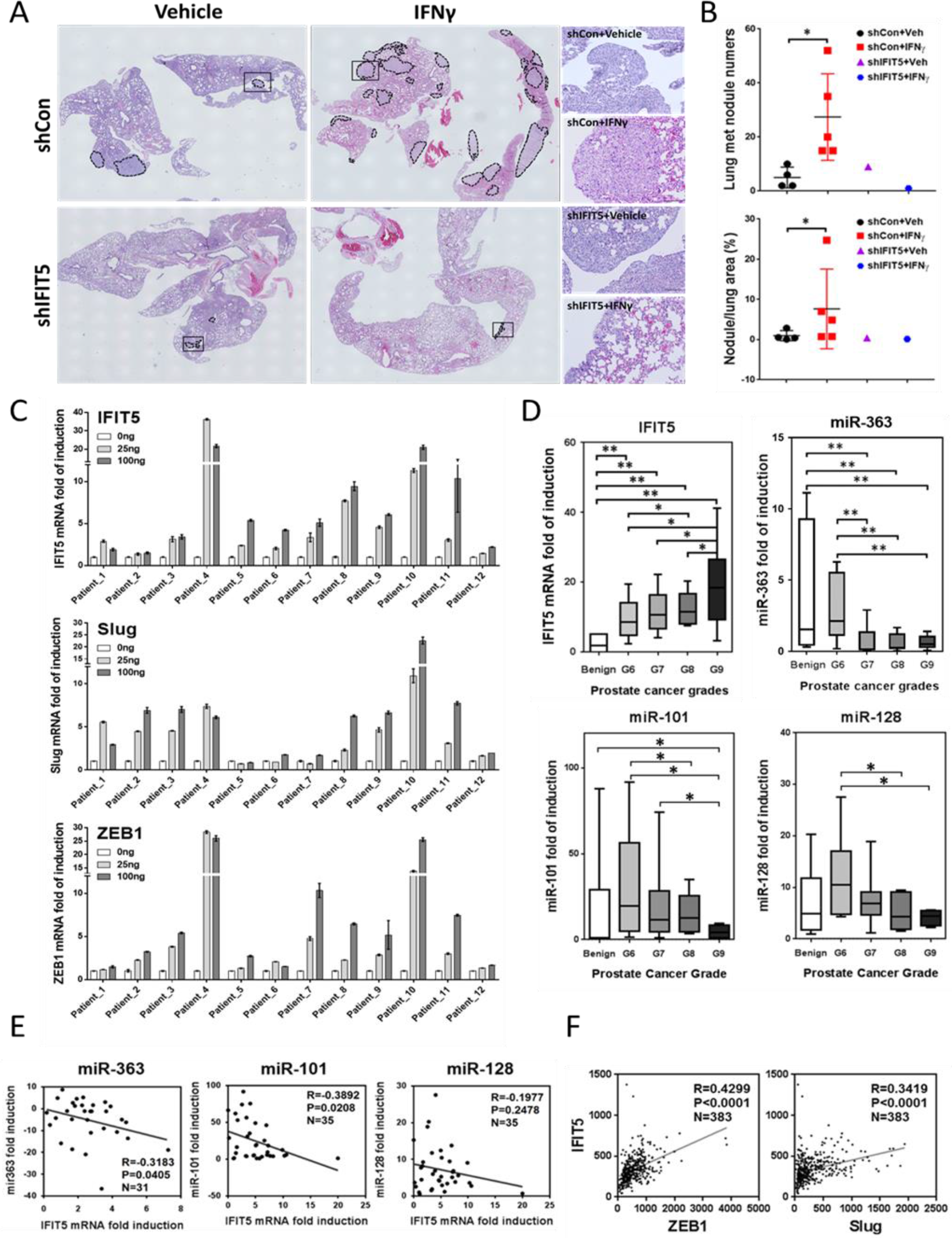
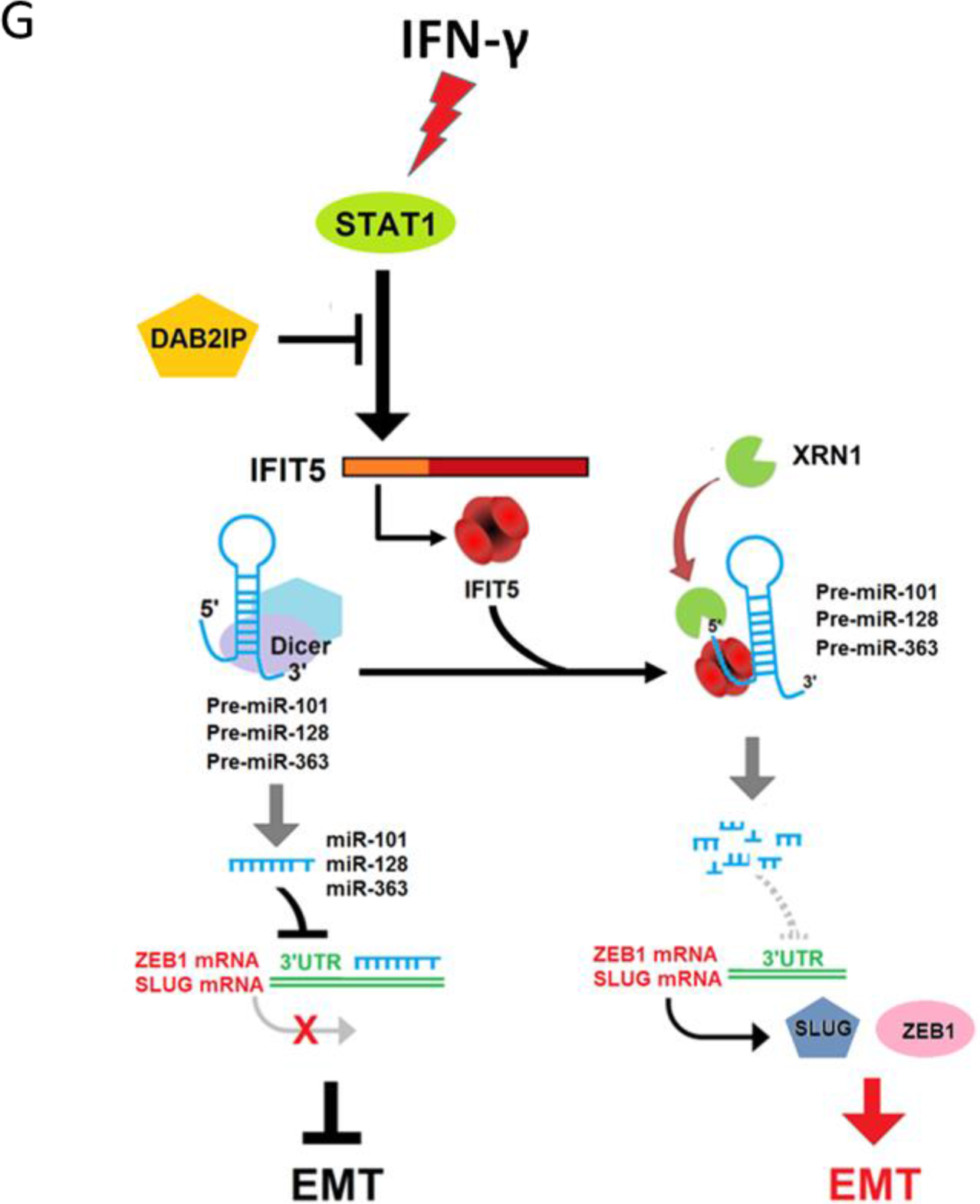
The effect of IFIT5-mediated miRNA degradation on EMT and its clinical correlation in PCa. (A) HE staining of lung tissue derived from mice receiving tail vein intravenous injection of IFIT5-knockdown PC3 cells (PC3-shIFIT5) pretreated with vehicle (Veh, PBS) or IFNγ (20ng/ml), compared to PC3 cells transfected with control vector (shCon). The black dotted line-circles region indicate the presence of metastatic nodules observed at lung parenchyma. Representative tumor nodules from each group are shown at right side panels (scale bar=100μm). (B) Comparison of tumor nodule numbers and comparative area ratio in the lung parenchyma among each group. (C) Induction level of IFIT5, Slug, and ZEB1 mRNA expression in *ex vivo* culture of human PCa specimens (N=12) treated with IFNγ (0, 25 and 100 ng/ml) for 48hrs. (D) Relative expression of IFIT5 mRNA and mature miR-363, miR-101 and miR-128 in human PCa specimens derived from different grades including benign (N=10), G6 (N=9), G7 (N=9), G8 (N=6) and G9 (N=7) (*p<0.05, **p<0.0001). (E) Clinical correlation of miR-363, miR-101 and miR-128 with IFIT5 mRNA expression in human PCa specimens graded from benign, G6 to G9. (F) Clinical correlation between IFIT5 and ZEB1 or Slug mRNA level in PCa from TCGA PCa dataset. (G) Schematic representing IFN induced IFIT5-mediated precursor microRNA degradation leading to EMT in cancer.

## DISCUSSION

A recent study using whole exome and whole transcriptome sequencing of patients with metastatic tumors demonstrated a strong correlation between cancer metastasis and the expression of interferon-induced genes or EMT. PCa is often found to have many different kinds of infiltrated immune cells such as macrophages, dendritic cells and tumor-infiltrating lymphocytes^23,24^. Instead of eliciting tumor immunity, these immune cells with secreting cytokines are capable of facilitating PCa development. For example, a study has demonstrated that fibroblast growth factor 11 (FGF11) released by the recruited CD4+ T cells can induce cell invasion by increasing matrix metalloproteinase 9 (MMP9) in PCa cells^25^. In addition, IL-4 produced from CD4+ T cells has shown to increase PCa cell survival and proliferation by activating the JNK signaling pathway in cancer cells^26^. Moreover, IL-17 secreted from T helper cells is capable of facilitating PCa invasiveness by increasing several EMT transcription factors and MMP7^27^. On the other hand, IFNγ, a type II interferon derived predominantly from CD4+/CD8+ lymphocytes and NK cell, is shown to have antitumor activities during innate immune response. Also, IFNγ has been used as a therapeutic agent exhibiting anti-proliferative^28^, anti-metastatic^29^, pro-apoptotic^30–33^ and anti-angiogenesis^34–37^ effects in various cancer types. However, several reports indicate that IFNγ could also facilitate tumor progression. For example, IFNγ can elicit CD4+ T-cell loss and impair secondary anti-tumor immune responses after initial immunotherapy using tumor-bearing mouse model ^38^. In colorectal carcinoma, IFNγ has been shown to facilitate the induction of indoleamine 2, 3 dioxygenase (IDO) that induces the production of kynurenines metabolites and impairs the function of surrounding T cells^39^. In addition to its role in immune modulation, blockade of IFNγ receptor (IFNGR) can inhibit peritoneal disseminated tumor growth of ovarian cancer^40^. Noticeably, serum IFNγ levels become elevated after radiotherapy in PCa patients^41^. Nevertheless the effect of IFNγ on the overall survival of PCa patients remains controversial^42^. In our study, we provide additional evidence that IFNγ and two other subtypes (Fig. 2F) are able to induce EMT, leading to cancer invasiveness via IFIT5-mediated turnover of tumor suppressor miRNAs (Fig. 6C and 7A-B). We also noticed that low concentration of IFNγ capable of inducing EMT exhibited no cytotoxicity (Fig. S9D). To strengthen the clinical evidence of IFN-induced EMT, we treated *ex vivo* PCa explants with IFNγ and demonstrated that IFNγ could induce similar elevations of IFIT5 and EMT transcriptional factors and suppression of miR-101, miR-128, and miR-363 (Fig. 8C and S9B). Taken together, these data show that IFNγ has a biphasic effect on tumor development. Nevertheless, the pro-tumorigenic effect of IFNγ at low concentration is expected to raise a concern for its application as an anti-tumor or immunotherapeutic agent.

Unlike other IFIT family proteins, IFIT5 is characterized as a monomeric protein that is capable of binding to viral RNA with 5’-triphosphate group^4^ as well as a broad spectrum of cellular RNA with either 5’-monophosphate or 5’-triphosphate group, including tRNA and other RNA polymerase III transcripts^6^. However, the interaction of IFIT5 with miRNA is largely unknown. Knowing that precursor miRNA shares a similar stem loop structure with tRNA and a precursor miRNA still retains 5’-monophosphate group after processing from its primary transcript, we are able to show that IFIT5 is capable of interacting with 5’-end of pre-miRNA molecules. After binding to pre-miRNA, IFIT5 recruits XRN1 to form unique miRNA turnover complex (Fig. 5). For the first time, we demonstrated that the specificity of miRNA recognition by IFIT5 is mainly determined by the 5’-end overhang structure of pre-miRNAs (Figs. 4, 6). Interestingly, these 3 tumor suppressor miRNAs (i.e., miR-101, miR-128 and miR-363) share similar 5’-end structure in their pre-miRNA and function in suppressing EMT despite of targeting different EMT transcriptional factors such as ZEB1 and Slug. To conclude, our study provides a new functional role of IFIT5 in miRNA biogenesis (Fig. 8G), particularly, a new understanding of differential regulation of cluster miRNAs.

Until now, the clinical correlation of IFIT5 in PCa is largely undetermined. In this study, we were able to demonstrate that the expression of IFIT5 is elevated in high-grade tumor and inverse correlation between IFIT5 and miR-101, −128 and −363 in PCa tumor specimens as well as from PCa TCGA database; this correlative relationship was not observed in other members of the miR-106a-363 cluster. In addition, a significant clinical correlation between IFIT5 and EMT transcription factors (ZEB1 or Slug) was observed from PCa TCGA dataset, which further validate the regulatory network of IFIT5-miRNAs-EMT. Taken together, data from clinical specimens provide a strong support for the key role of IFIT5 in EMT in human PCa by regulating the turnover of tumor suppressor miRNAs targeting EMT transcription factors, which could contribute to the design of miRNA therapy. Also, with respect to the clinical correlation of IFIT5 and its mechanism of action in PCa and renal cancer cells, IFIT5 can be a potential therapeutic target.

## MATERIALS AND METHODS

### Cell lines, Clinical specimens, and Plasmid constructs

Stable DAB2IP-KD and control (Con) prostatic cell lines were generated from RWPE-1, PC-3 and LAPC4 cell lines using shRNA^16^. Stable IFIT5-knockdown (shIFIT5) and control (shCon) prostatic cell lines were generated from PC3, LAPC4-KD, RWPE1-KD and C4-2Neo cell lines using pLKO-shIFIT5 from Academia Sinica, Taipei, Taiwan. Stable IFIT5-overexpressing (IFIT5) and control (Vec) prostatic cell lines were generated from LAPC4-Con, RWPE1-Con and C4-2D2 cell lines using pcDNA3.1-3XFlag-IFIT5 plasmid from Dr. Collins. LAPC4 derived from PCa patients with lymph node metastasis was maintained in Iscove Dulbecco’s Modified Eagle’s Medium (DMEM) (Invitrogen) containing 10% fetal bovine serum (FBS). RWPE-1 derived from normal prostate epithelial cells immortalized with human papillomavirus 18 was maintained in Keratinocyte medium (Invitrogen) containing 10% FBS. The androgen-sensitive LNCaP cell line derived from PCa patients with lymph node metastasis was maintained in RPMI-1640 medium (Invitrogen) containing 10% FBS. C4-2 and PC-3, an androgen-independent line, were maintained in RPMI-1640 medium containing 10% FBS. Renal cancer 786O cell lines were maintained in RPMI-1640 medium (Invitrogen) containing 10% FBS, whereas ovarian carcinoma (HeLa) and kidney cell (293T) cell lines are maintained in DMEM (Invitrogen) containing 10% FBS.

A total of 41 PCa specimens obtained from UT Southwestern Tissue Bank. All the specimens were collected from 6-mm core punch from radical prostectomy and examined by pathologist to determine tumor grade then subjected to RNA extraction. The Institutional Review Board of UT Southwestern approved the tissue procurement protocol for this study, and appropriate informed consent was obtained from all patients.

All the plasmid constructs are described in Supplemental information.

### Cell transfection

Cells (2.5×10^5^) were seeded in 60-mm dish at 60-70% confluence before transfection. According to manufacturer’s protocol, transfection of plasmids was using either Xfect Reagent (Clontech) or EZ Plex transfection reagent (EZPLEX), and transfection of XRN1 or IFIT5 siRNA was using Lipofectamine^®^ RNAiMAX reagent (Life Technology). Transient transfection was carried out 48 hrs post-transfection to harvest cell for further analyses. In addition, the stable clones (CL) were established after 2 weeks in the antibiotic selective medium.

### RNA isolation and quantitative real-time RT-PCR (qRT-PCR)

Small and large RNA were isolated and purified using mirVana miRNA Isolation Kit (Life Technologies). Small RNA (250 ng) was subjected to miScript II RT kit (QIAGEN) then 2.5 μl cDNA was applied to a 25-μL reaction volume using miScript SYBR Green PCR kit (QIAGEN) in iCycler thermal cycler (Bio-Rad). All primer sequences are listed in Table S3. The relative expression levels of precursor and matured miRNAs from each sample were determined by normalizing to SNORD95 small RNA. Large RNA (2 μg) was subjected to SuperScript VILO cDNA synthesis kit (Invitrogen) then 2.5 μl cDNA was applied to 25-μl reaction volume using SYBR Green ER qPCR SuperMix (Invitrogen). The relative expression levels of DAB2IP, IFIT5, E-cadherin, and Vimentin, ZEB1 and Slug/SNAI2 mRNA from each sample were determined by normalizing to 18S mRNA.

### Western blot analysis

Cells were washed with PBS and lysed in lysis buffer [50mMTris-HCl (pH 7.5), 150 mM NaCl, 0.1% Triton X-100, 1 mM sodium orthovanadate, 1 mM sodium fluoride, 1 mM sodium pyrophosphate, 10 mg/mL, aprotinin, 10 mg/mL leupeptin, 2 mM phenylmethylsulfonyl fluoride, and 1 mM EDTA] for 60 mins on ice. Cell lysates were spin down at 20,000 xg for 20 mins at 4°C. Protein extracts were subjected to SDS-PAGE using Bolt 4-12% Bis-Tris Plus gel (Invitrogen), and transferred to nitrocellulose membrane using Trans-Blot Turbo Transfer system (BIORAD). Membranes were incubated with primary antibodies against DAB2IP (AbCam), E-Cadherin (BD Transduction Laboratory), Vimentin (Sigma), ZEB1 (Cell signaling Technology), Slug/SNAI2 (Cell signaling Technology), XRN1 (Santa Cruz Biotechnology), IFIT5 (ProteinTech) and HRP-Flag (Sigma) at 4 °C for 16-18 hrs, and horseradish peroxidase-conjugated secondary antibodies at room temperature for 1.5 hrs. Results were visualized with ECL chemiluminescent detection system (Pierce ThermoScientific). The relative protein expression level in each sample was normalized to actin or GAPDH.

### Immunoprecipitation (IP) assay

Cells were harvested and protein lysates were prepared freshly before performing IP assay. Flag antibody (Sigma) or XRN1 antibody (AbCam) was incubated with 50 μl of Dynabeads^®^ protein G (Novex, Life Technology) at room temperature for 15 mins. Subsequently, total 300 μg of protein lysate was incubated with the Dynabead-conjugated antibody at 4°C for 16-18 hrs. After washing, the elutes were subjected to SDS-PAGE using Bolt 4-12% Bis-Tris Plus gel (Invitrogen), and transferred to a nitrocellulose membrane using Trans-Blot Turbo Transfer system (BIORAD). Membranes were then subjected to western blot probed with XRN1 or Flag antibody.

### Luciferase reporter assay

Cells (8×10^4^) were seeded onto 12-well plates at 75% confluence before transfection. pCMV-miR-363 plasmid was co-transfected with psiCHECK2-Slug3’UTR-WT or psiCHECK2-Slug3’UTR-Mut^363^ plasmid into LAPC4-KD or RWPE-1-KD cells using Xfect Reagent (Clontech). Cells were harvested and lysed with Passive Lysis buffer (Promega) at 48 hrs after transfection. Luciferase activity was measured using the Dual-luciferase reporter assay (Promega) on the Veritas Microplate Luminometer (Turner Biosystems). Results were expressed as the relative light unit (RLU) by normalizing the firefly luciferase activity with Renilla luciferase activity. Each experiment was performed in triplicates.

### *In vitro* transcription of precursor miRNA

The PCR-amplified DNA fragment of T7-precursor-miRNAs was separated by 2% agarose gel electrophoresis and purified using Mermaid SPIN kit (MP Biomedicals), then subjected to *in vitro* transcription assay using T7 High Yield RNA synthesis kit (New England Biolabs). DNA template (750 ng) was mixed with T7 High Yield 10X buffer, NTP mixture (ATP, GTP, CTP and UTP), and T7 RNA polymerase then incubated at 37°C for 16 hrs. The precursor miRNA molecules was first treated with DNase I for 15 mins at 25°C and purified by acid phenol-chloroform extraction and ethanol precipitation at −20°C for 1hr. The molecular size and sequence of each purified precursor miRNA was confirmed by gel electrophoresis using 15% TBE-Urea gel and qRT-PCR respectively.

### RNA pull-down assay

The *in vitro* transcribed precursor miRNA was subjected to RNA pull down assay using Pierce Magnetic RNA-Protein Pull-Down Kit (ThermoScientific). An approximate 100 pmol of precursor miRNA were incubated with 10X RNA Ligase reaction buffer, RNase inhibitor, Biotinylated Cytidine Bisphosphate, and T4 RNA ligase at 16°C for 16 hrs. The biotinylated precursor miRNA was then purified and incubated with streptavidin magnetic beads for 30 mins at room temperature. Whole cell lysates were freshly prepared immediately before RNA pull-down assay and 200 μg of protein extract was mixed with the biotinylated precursor miRNA conjugated to streptavidin magnetic beads and incubated at 4°C for 1 hr. The magnetic beads were washed 4 times before elution. Proteins associated with precursor miRNA were eluted and subjected to SDS-PAGE using Bolt 4-12% Bis-Tris Plus gel. For <u>unveiling pre-miR-363-associated protein candidates, gel bands were stained with Coomassie blue and subjected to LC-MS/MS analysis. For identifying the interaction between mutant pre-miR-363s (SS6Mut, DSMut) and IFIT5 or microRNA processing machinery, the gel was transferred to nitrocellulose membrane using Trans-Blot Turbo Transfer system (BIORAD). Membranes were incubated with primary antibodies against IFIT5, Dicer or Drosha proteins.</u>

### *In vitro* pre-miRNA degradation assay

The *in vitro* transcribed precursor miRNAs were incubated with immunoprecipitated IFIT5 or XRN1 protein, as well as recombinant rIFIT5 protein and/or XRN1 enzyme (New England Biolab) in the elution buffer at 37°C on a thermomixer (Eppendorf), then the RNA-containing buffer were collected at indicated time points and subjected to 15% TBE-Urea gel electrophoresis. To quantify the degradation of precursor miRNA, the 15% TBE-Urea gel was then stained with GelRed− Nucleic Acid Gel Stains (VWR) and visualized under UV light in the Alpha Imager devise (Protein Simple). The RNA bands were quantified by Multiplex band analysis (AlphaView Software) and the rate of degradation was calculated from each time point normalized to time zero.

### *In vitro* migration and invasion assay

Cells (1×10^5^ or 4×10^4^) in the serum-free medium were plated on the upper chamber (8−)m pore size) of Transwell (Corning) with or without Matrigel coating for invasion or migration assay, respectively, while lower chamber contained medium supplemented with 10% FBS. After 5 days, cells that had transmigrated to the lower chamber were fixed by 4% paraformaldehyde, stained and visualized under microscope. Quantification of migratory cells was carried out with crystal violet staining and measurement at OD_555nm_. Each experiment was performed in triplicates.

### Mouse intravenous injection

Stable clones of PC3-shCon or-shIFIT5 were infected with luciferase lentivirus. One million PC3 cells pre-treated with Vehicle (PBS) or IFNγ (20 ng/ml, 48 hrs) were resuspended in 50μl PBS and then injected into the tail vein of male SCID mice, followed by intravenous injection of IFNγ (5 ng/ml,) weekly for 4 weeks. At 8^th^ week post-injection, lung metastasis of PC3 tumor was observed by bioluminescent imaging (BLI) using IVIS^®^ system then lungs were excised, fixed in 10% formalin, paraffin-embedded and stained with HE for pathological identification of tumor nodules presented in the lung parenchyma.

### *Ex vivo* culture of patient-derived PCa explants

Following informed consent, a total of 12 fresh PCa tissues were obtained from men undergoing radical prostatectomy at the hospitals of the University of Texas Southwestern Medical Center (Dallas, TX). The *ex vivo* culture was performed as previously described^21^. Briefly, fresh PCa tissue was dissected into 1mm^3^ cube and placed on a Gelatin sponge (Novartis, East Hanover, NJ) bathed in RPMI-1640 media supplemented with 10% heat-inactivated FBS, 100units/mL penicillin-streptomycin, 0.01 mg/mL hydrocortisone and 0.01 mg/mL insulin (Sigma). In addition, to the media, was added either vehicle, IFNγ (25 ng/ml) or IFNγ (100 ng/ml). Tissues were cultured at 37 °C for 48 hrs then snap-freeze in liquid nitrogen for RNA purification.

### Statistics analysis

Statistics analyses were performed by using GraphPad Prism software. Statistical significance was evaluated using Student t-test. P<0.05 was considered a significant difference between compared groups and marked with an asterisk. The statistical association between miR-363 and IFIT5 expression among different grades of human PCa was evaluated with regression correlation analysis.

## ACKNOWLEDGMENTS

We thank Dr. Collins (University of California, Berkeley) for providing IFIT5 cDNA constructs, Dr. Dong (Emory University, Atlanta) for providing the psiCHECK2-Slug3’UTR plasmid. Drs. Kou-Juey Wu (China Medical University, Taichung, Taiwan) and Dr. Vimal Selvaraj (Cornell University, Ithaca) for the helpful discussion. This work was supported by grants from the United States Army (W81XWH-11-1-0491 and W81XWH-16-1-0474 to JTH) and (W81XWH-14-1-0249 to UGL), and the Ministry of Science and Technology in Taiwan (MOST103-2911-I-005-507 to HL)

## SUPPLEMENTAL FIGURES AND FIGURE LEGENDS

**Figure S1.**
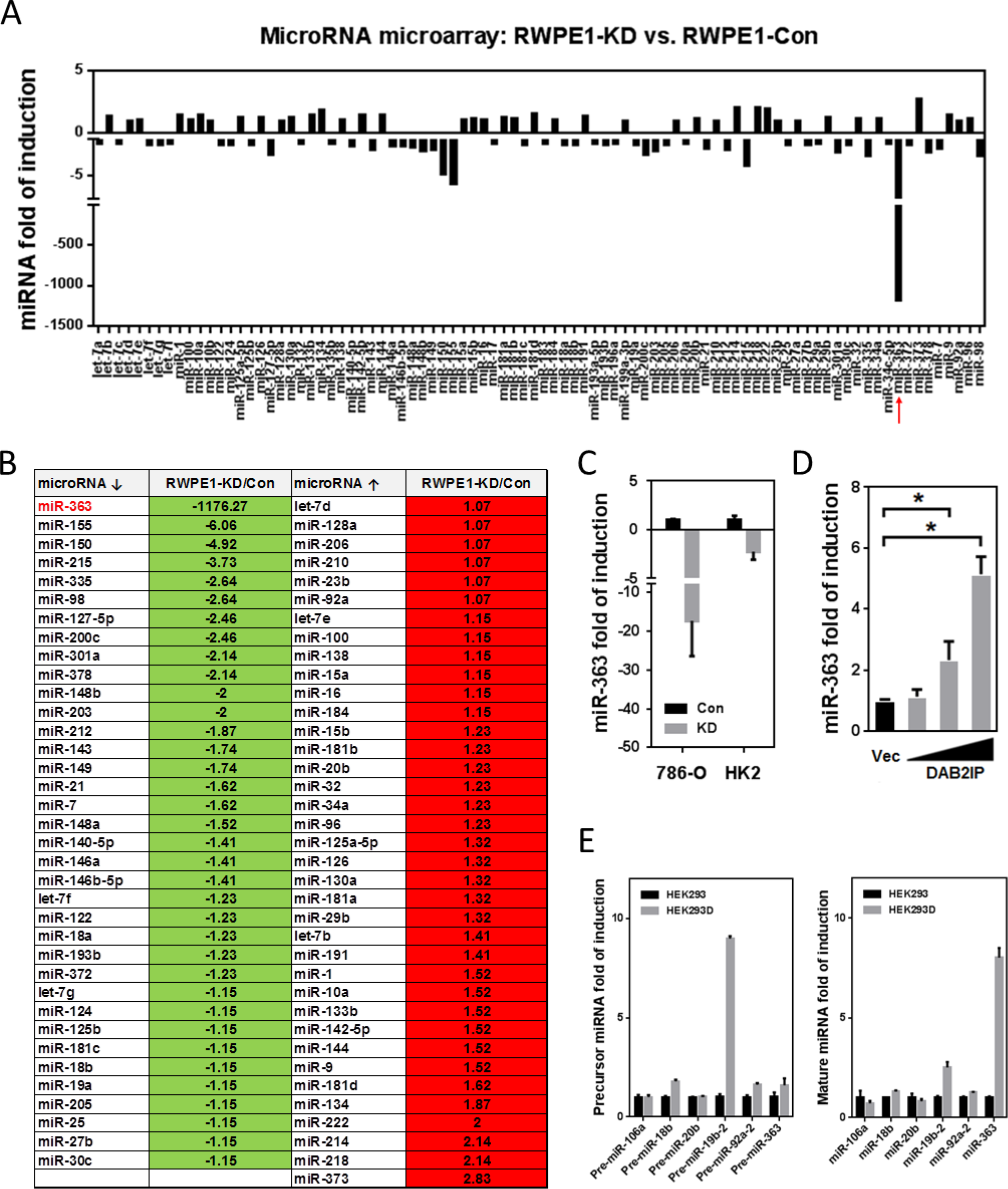
(A) Profile of miRNA expression in RWPE1-KD cells compared with RWPE1-Con cells. (B) Relative fold change of each miRNA from microarray screening (green and red indicate decreased and increased fold change in RWPE1-KD cells after normalizing with RWPE1-Con cells, respectively). (C) The expression levels of mature miR-363 in DAB2IP- KD renal cell lines (786-O and HK2) after normalizing with the vector control (Con). (D) Induction of mature miR-363 by ectopic expression of DAB2IP in HEK293 cells. (E) Expression levels of precursor and mature miRNAs (miR-106a, iR-18b, miR-20b, miR-19b- 2, miR-92a-2 and miR-363) in DAB2IP-expressing HEK293D cells after normalizing with HEK293 cells.

**Figure S2.**
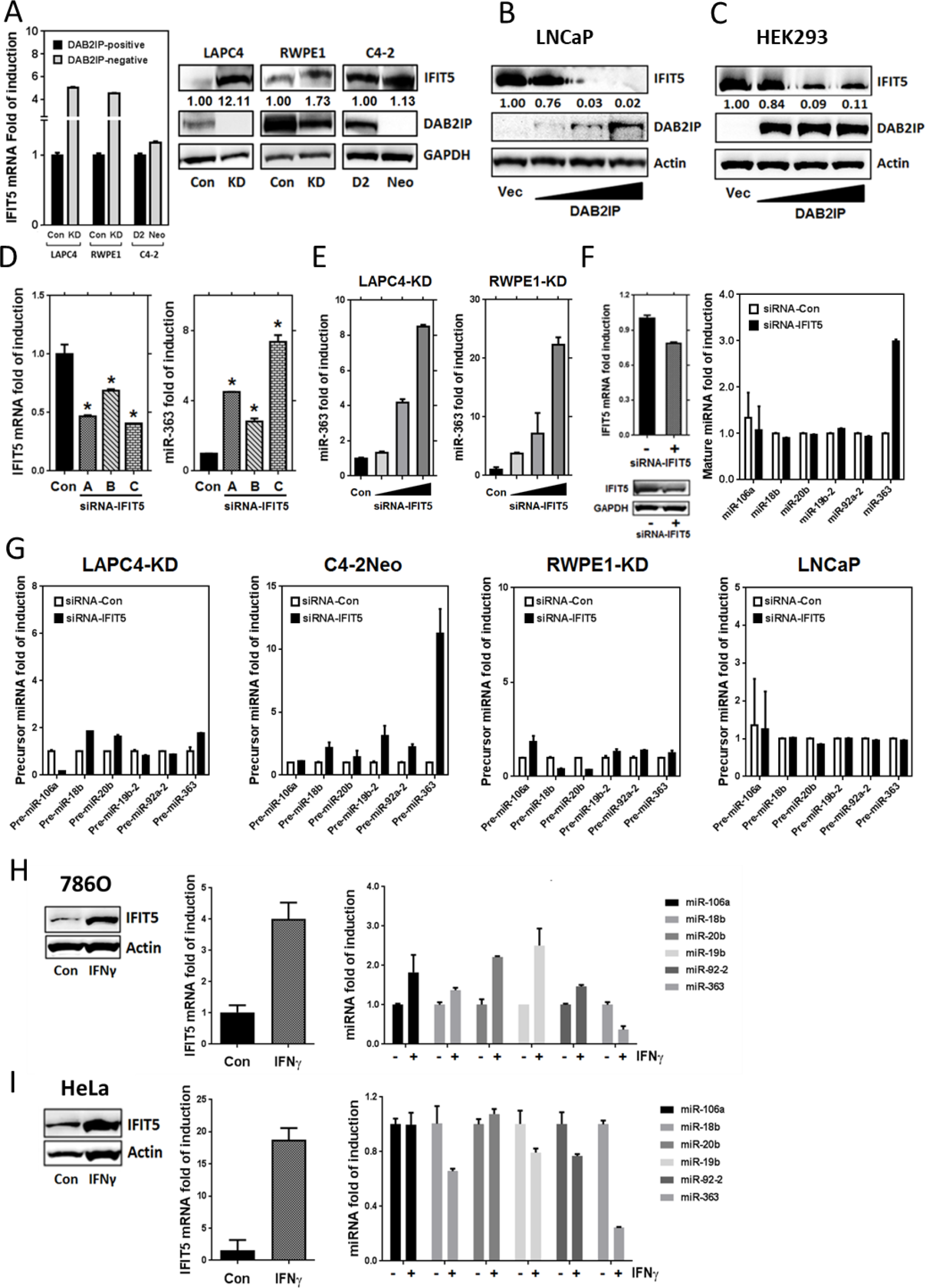
(A) Relative change of IFIT5 mRNA and protein level in pairs of DAB2IP-positive and -negative prostate cells (LAPC4, RWPE1 and C4-2). (B-C) Suppression of IFIT5 protein expression by ectopic expression of DAB2IP in LNCaP and HEK293 cell lines after normalizing with the control vector (Vec). (D) Induction of mature miR-363 in LAPC4-KD cells transfected with IFIT5 siRNAs compared with control siRNA (Con). (E) Induction of mature miR-363 in LAPC4-KD and RWPE1-KD cells by IFIT5 siRNA knockdown after normalizing with the control siRNA (Con). (F) Expression levels of mature miRNAs (miR- 106a, miR-18b, miR-20b, miR-19b-2, miR-92a-2 and miR-363) in IFIT5-KD (siRNA-IFIT5/+) LNCaP cells compared with the control siRNA (siRNA-Con/−). (G) Expression levels of precursor miRNAs (miR-106a, miR-18b, miR-20b, miR-19b-2, miR-92a-2 and miR-363) in IFIT5-KD (siRNA-IFIT5) LAPC4-KD, C4-2Neo, RWPE1-KD and LNCaP cells after normalizing with the control siRNA (siRNA-Con). (H-I) Left: Induction of IFIT5 protein in 786O and HeLa cells treated with IFNγ (10ng/ml) for 48 hrs. Right panel: Expression levels of mature miRNAs (miR-106a, miR-18b, miR-20b, miR-19b-2, miR-92a-2 and miR-363) in 786O and HeLa cells treated with IFNγ (10ng/ml) for 48 hrs. (J) Induction of IFIT1 and IFIT5 protein in RWPE1 or PC3 cells treated with IFNγ (0, 5 and10ng/ml) for 48 hrs.

**Figure S3.**
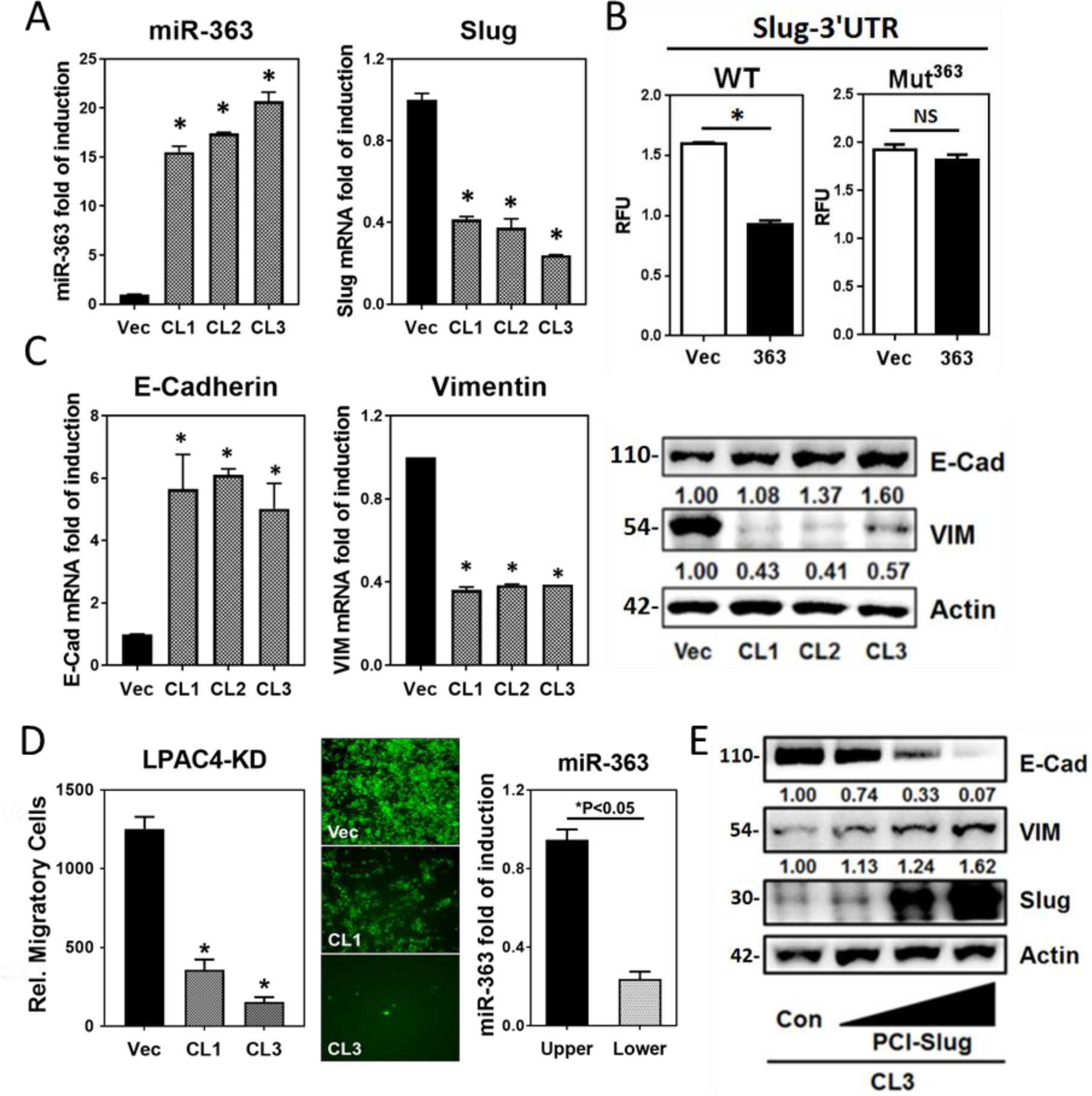
Reduction of Slug/SNAI2 mRNA levels in miR-363-expressing LAPC4-KD cells after normalizing with the control vector (Vec). (*p<0.05, CL: miR-363 expressing stable clone) (B) Luciferase reporter activities in LAPC4-KD cells co-transfected with siCHECK2-Slug-WT 3’UTR or -Slug Mut363 3’UTR and pCMV-miR363 or control vector. (RFU=Renilla to Firefly luciferase activity, each bar represents mean ± SD of four replicated experiments. * p<0.05) (C) Expression levels of E-cadherin and Vimentin mRNA and protein level in miR-363-expressing LAPC4-KD cell. (D) Left and middle: The effect of miR-363 on cell migration of GFP–expressing LAPC4-KD cells. Migrated GFP-positive cells were observed under microscope and stained with crystal violet and quantified at O.D. 555nm. (Each bar represents mean ± SD of three replicated experiments. *p<0.05). Right: miR-363 expression level in LAPC4-KD cells migrated to the lower chamber, compared to cells stay at upper chamber. (CL: miR-363 expressing stable clone of LAPC4-KD cells) (E) The effect of Slug on the expression levels of E-cadherin and Vimentin mRNA in miR-363-expressing LAPC4-KD cells (CL3) after normalizing with the control vector (Con). (*P<0.05).

**Figure S4.**
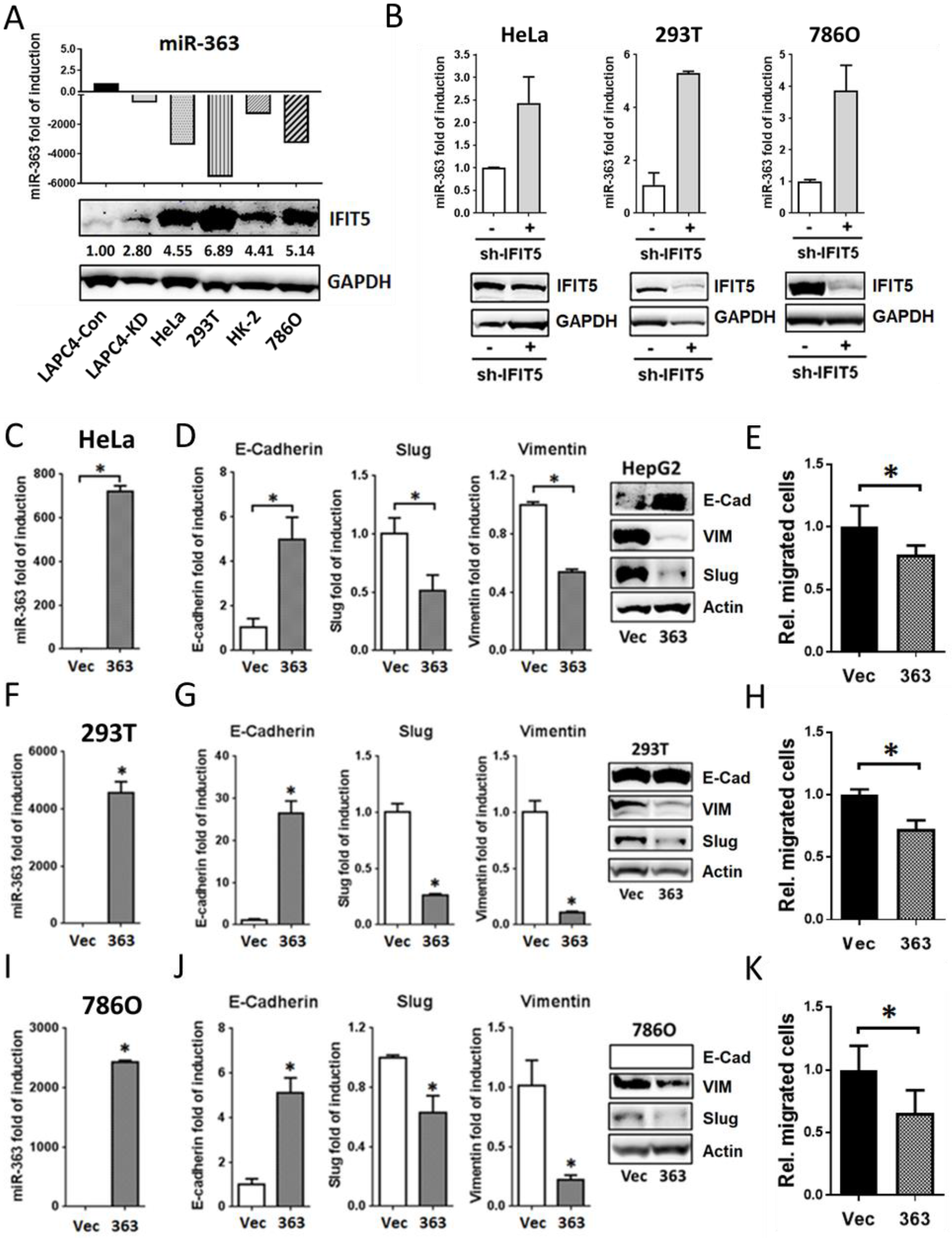
(A) The expression level of miR-363 and IFIT5 protein among prostate (LAPC4), ovarian (HeLa) and kidney (293T, HK2 and 786O) cancer cell lines. (B) Induction of mature miR-363 levels in IFIT5-shRNA knockdown (shIFIT5, +) HeLa, 293T and 786O cell lines, compared to the control shRNA (−). (C, F, I) Overexpression of miR-363 in HeLa, 293T and 786O cell lines. (D, G, J) The impact of miR-363 on the mRNA and protein level of E-cadherin, Slug and Vimentin in HeLa, 293T and 786O cell lines transfected with pCMV-miR- Dell migration of miR-363 expressing HeLa, 293T and 786O cells (363), compared to the cells with control vector (Vec).

**Figure S5.**
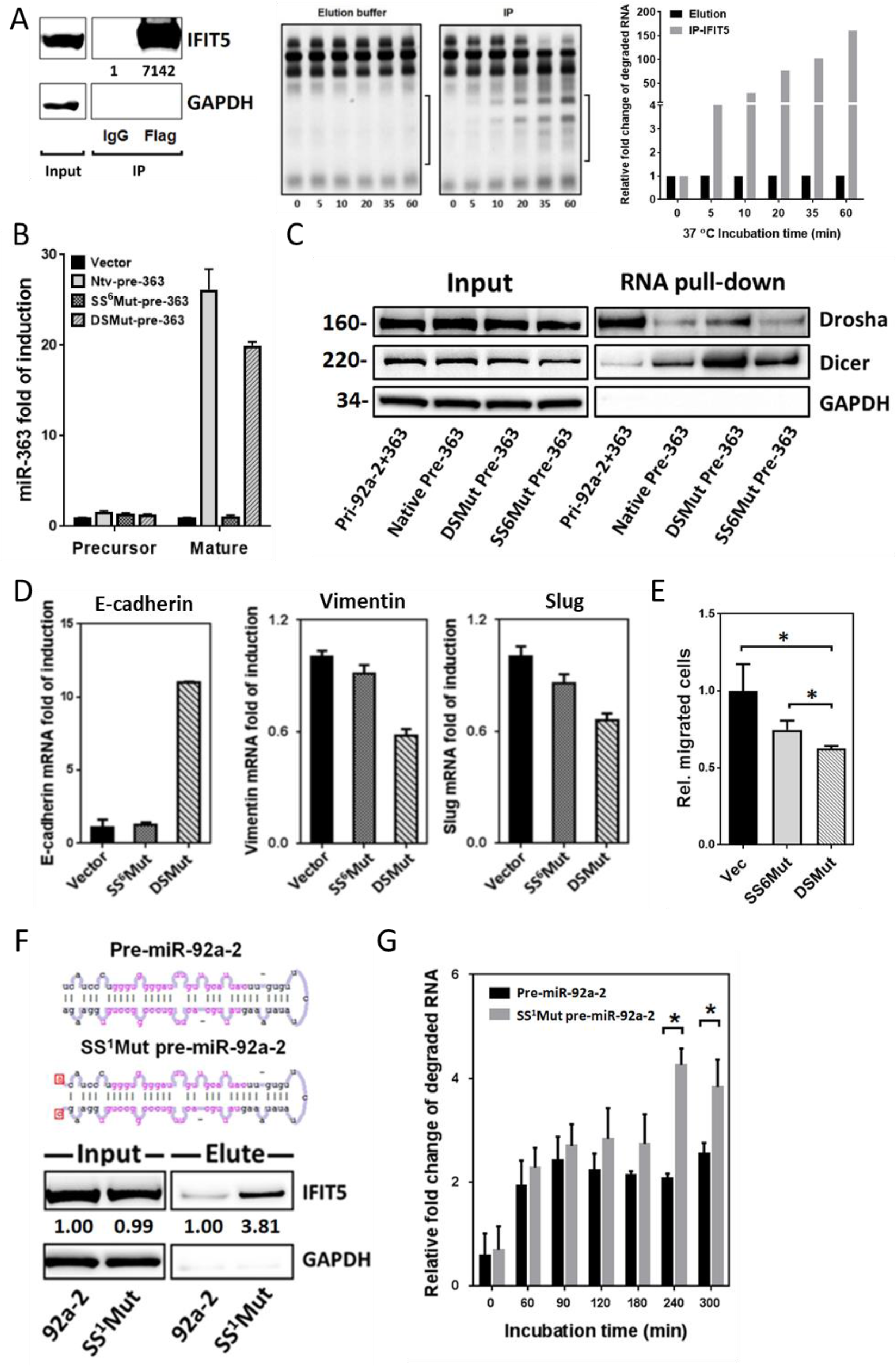
(A) Left: Immunoprecipitation (IP) of ectopic expressed IFIT5 protein subjected to in vitro RNA degradation assay. Middle panel: Gel electrophoresis of Native pre-miR-363 fragments after incubation with IFIT5 protein (IP) or elution buffer at 37°C. Right: Time-dependent change of degraded pre-miR-363 fragments (bracket) after normalized with 0 min. (B) Expression levels of precursor and mature miR-363 in RWPE1-KD cells transfected with native, SS6Mut or DSMut pre-miR-363 plasmids for 24 hrs and normalized with the control vector (Vec). (C) Interaction between Drosha or Dicer protein and primary miR-92a-2-miR- 363 (pri-92a-2+363, positive control for Drosha binding), native, SS6Mut or DSMut pre-miR-363 RNA molecules using RNA pull down assay. (D) The effect of SS6Mut and DSMut pre-miR-363 on the expression levels of mature E-cadherin, Vimentin and Slug mRNA in LAPC4-KD cells after normalizing with the control vector. (E) The effect of SS6Mut or DSMut pre-miR-363 on cell migration in LAPC4-KD cells. Migrated cells were stained with crystal violet and quantified at OD 555nm. Each bar represents mean ± SD of three replicated experiments. (* p<0.05) (F) Upper panel: predicted structure and sequence of pre-miR-92a-2. Middle panel: mutation of nucleotides (red box) for generating single nucleotide overhanging structure of pre-miR-92a-2 (SS1Mut). Lower panel: interaction between IFIT5 protein and pre-miR-92a-2 or SS1Mut pre-miR-92a-2 RNA molecules using RNA pull down assay. (G) Time-dependent change of degraded pre-miR-92a-2 and SS1Mut pre-miR-92a-2 after incubating with IFIT5 protein complex at 37°C normalized with 0 min. (*P<0.05).

**Figure S6.**
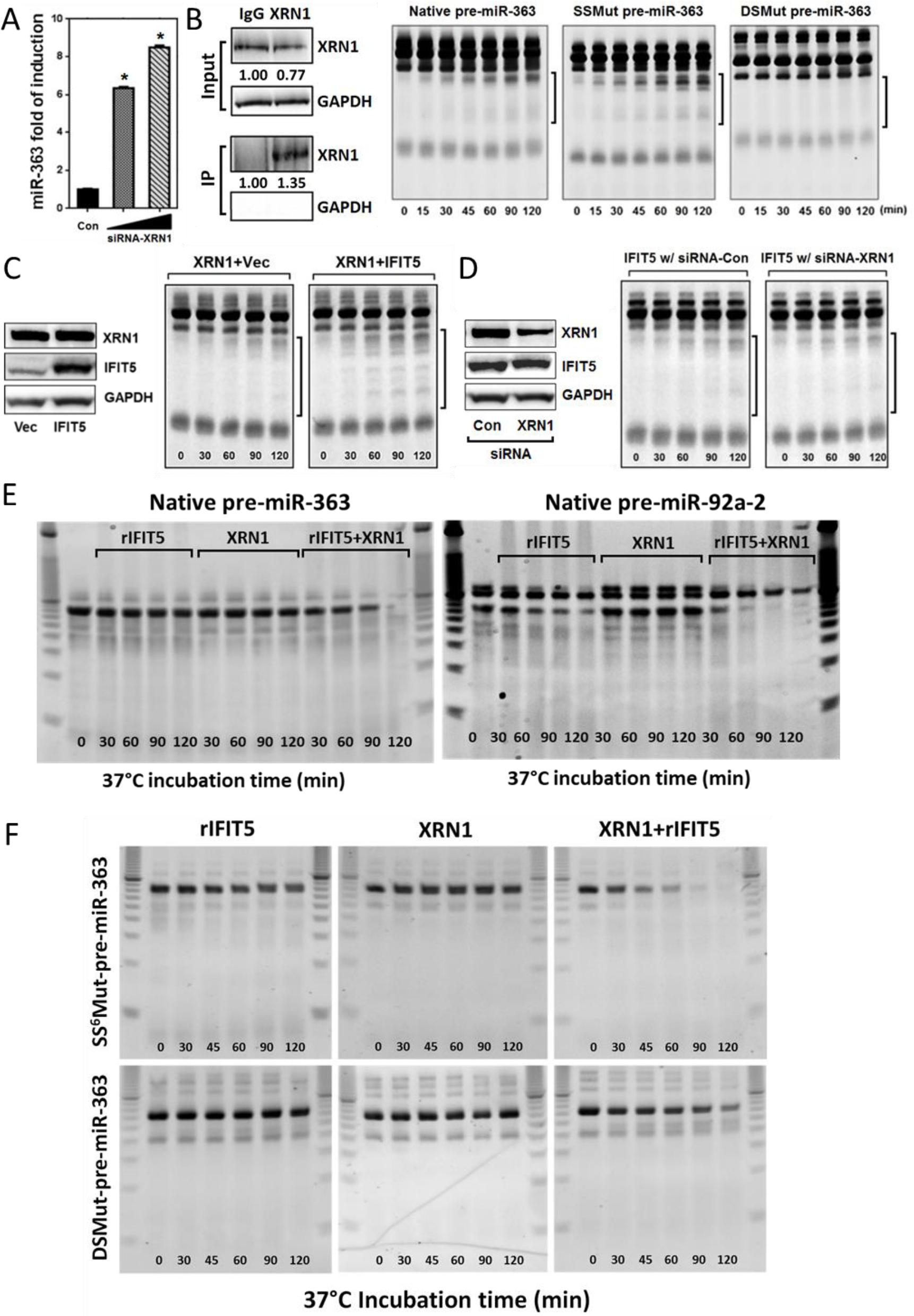
(A) Induction of miR-363 expression in LAPC4-KD cells transfected with XRN1 siRNA-B and compared with the control siRNA (Con). (B) Left panel: IP of endogenous XRN1 protein subjected to in vitro RNA degradation assay. Right panel: Time-dependent change of degraded native, SS6Mut and DSMut pre-miR-363 RNA molecules (bracket) after incubation with IP-XRN1 protein at 37°C, each time point was normalized with 0 min. (C) Left panel: Western blot from Vector or IFIT5-transfected LAPC4-Con cells. Right panel: Time-dependent change of degraded SS6Mut pre-miR-363 fragments in the presence of XRN1 alone (XRN1+Vector) or XRN1-IFIT5 complex (XRN1+IFIT5) at 37°C. (D) Left panel: western blot from IFIT5-expressing LAPC4-Con cells transfected with control or XRN1 siRNA. Right panel: Time-dependent change of degraded SS6Mut pre-miR-363 fragments in the presence of XRN1-IFIT5 complex derived from IFIT5 w/ siRNA-Con or IFIT5 w/ siRNA-XRN1 at 37°C. (E) In vitro RNA degradation of native pre-miR-363 or pre-miR-92a-2 after incubation with recombinant IFIT5 protein (rIFIT5), XRN1 enzyme (XRN1) or combination of XRN1 and rIFIT5 at 37°C for 0, 30, 60, 90 and 120 min. (F) In vitro RNA degradation of SS6Mut- or DSMut-pre-miR-363 after incubation with rIFIT5, XRN1 or combination of XRN1 and rIFIT5 at 37°C for 0, 30, 45, 60, 90, and 120 min.

**Figure S7.**
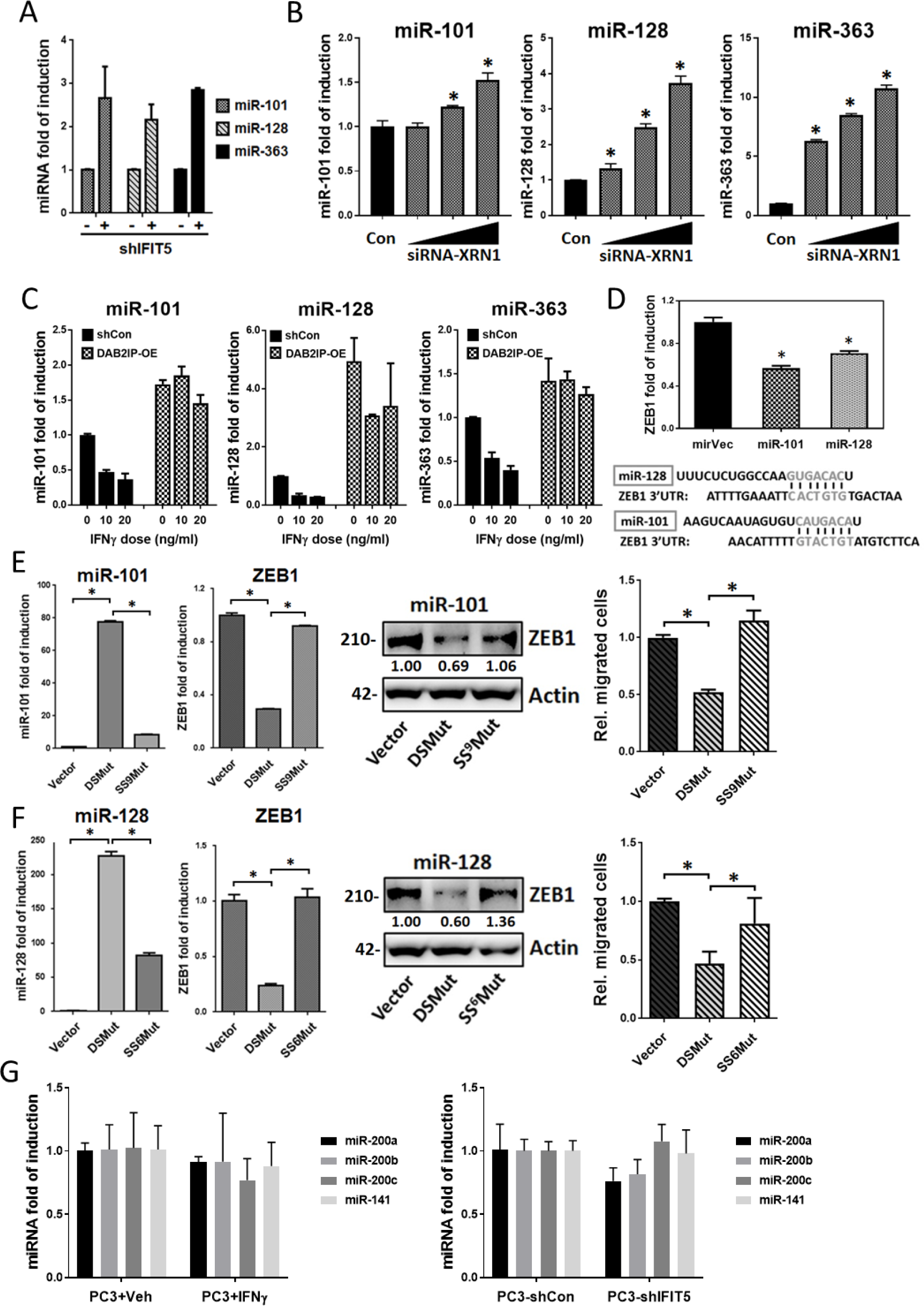
(A) Expression level of mature miR-101, miR-128 and miR-363 in IFIT5-shRNA knockdown LAPC4-KD. (B) Dose-dependent recovery of mature miR-363, miR-101 and miR-128 expression in IFIT5-expressing LAPC4-KD cells transfected with XRN1 siRNA after normalizing with the control vector (Con: control siRNA, *p<0.05).(C) Expression level of mature miR-101, miR-128 and miR-363 in DAB2IP-overexpressing PC3 cell lines treated with IFNγ (0, 10 and 20ng/ml) for 48hrs (*P<0.05). (D) Upper: Expression of ZEB1 mRNA level in PC3 cells overexpressed with miR-101 or miR-128, compared to vector control (miRVec). Lower: Matched sequence paired between the seed region of miR-101 or miR-128 and the 3’UTR of ZEB1 mRNA (E) Left and Middle panel: The effect of DSMut and SS9Mut pre-miR-101 on the expression levels of mature miR-101 and ZEB1 mRNA level in LAPC4-KD cells after normalizing with the control vector. Right panel: The effect of DSMut and SS9Mut pre-miR-101 on cell migration in LAPC4-KD cells. Migrated cells were stained with crystal violet and quantified at OD 555nm. Each bar represents mean ± SD of three replicated experiments. (* p<0.05, NS=no significance). (F) Left and Middle panel: The effect of DSMut and SS6Mut pre-miR-128 on the expression levels of mature miR-128 and ZEB1 mRNA level in LAPC4-KD cells after normalizing with the control vector. Right panel: The effect of DSMut and SS6Mut pre-miR-128 on cell migration in LAPC4-KD cells. Migrated cells were stained with crystal violet and quantified at OD 555nm. Each bar represents mean ± SD of three replicated experiments. (* p<0.05, NS=no significance) (G) Expression of miR- 200 family members in PC3 cells treated with IFNγ (Left panel) or knockdown with IFIT5 (right panel).

**Figure S8.**
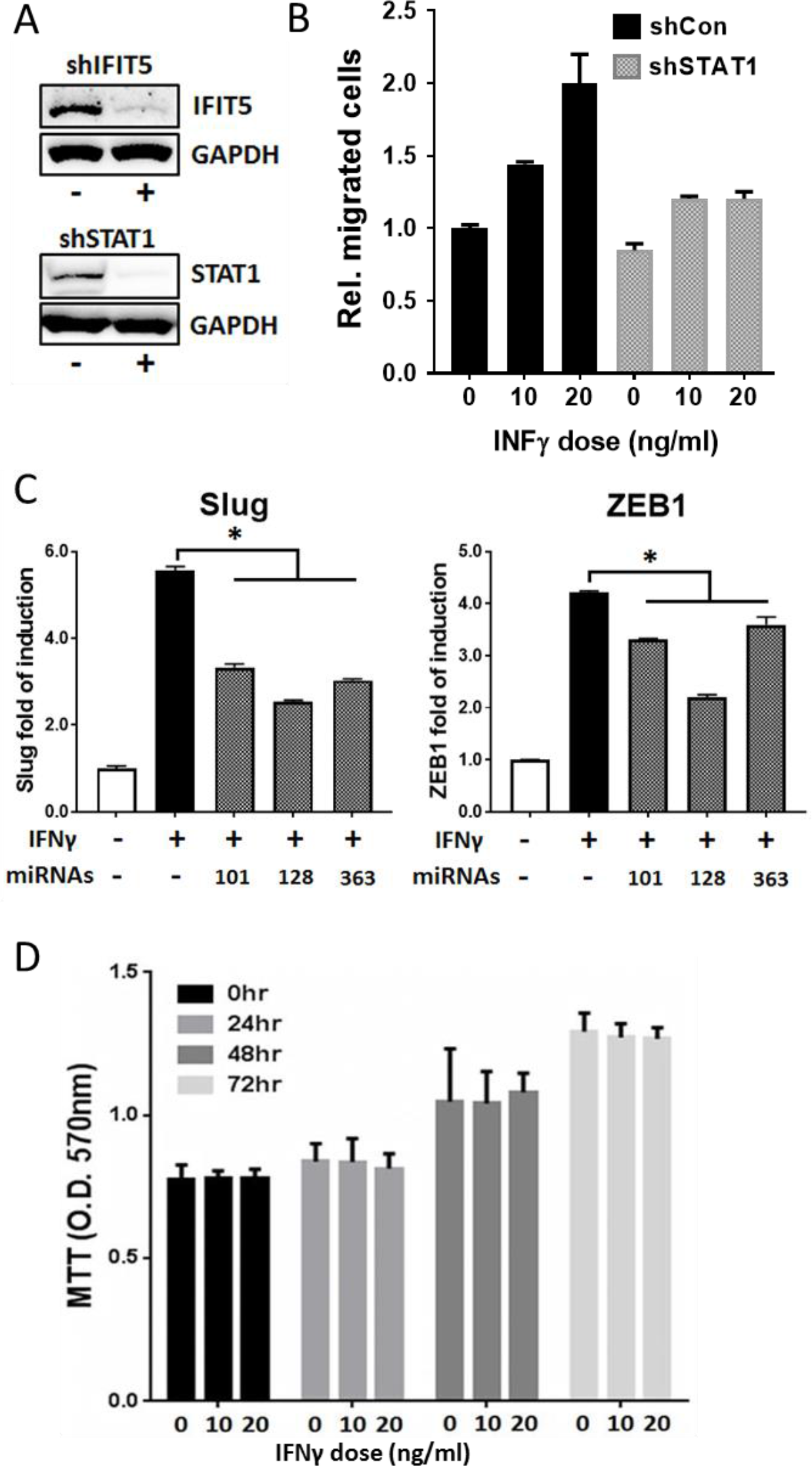
(A) Western blot demonstrating the knockdown of IFIT5 or STAT1 (+) in PC3 cells, compared to control shRNA (−). (B) Transwell migration of STAT1-shRNA knockdown (shSTAT1) PC3 cells after 48 hrs treatment of IFNγ, compared to shCon. Migrated cells were stained with crystal violet and quantified at OD 555nm. Each bar represents mean ± SD of three replicated experiments. (* p<0.05, NS=no significance) (C) IFNγ-induced upregulation of Slug and ZEB1 mRNA level in miR-101, miR-128 or miR-363-overexpressed PC3 cells. (D) Cell proliferation rate of PC3 cells treated with different dose of IFNγ for 24, 48 and 72 hrs.

**Figure S9.**
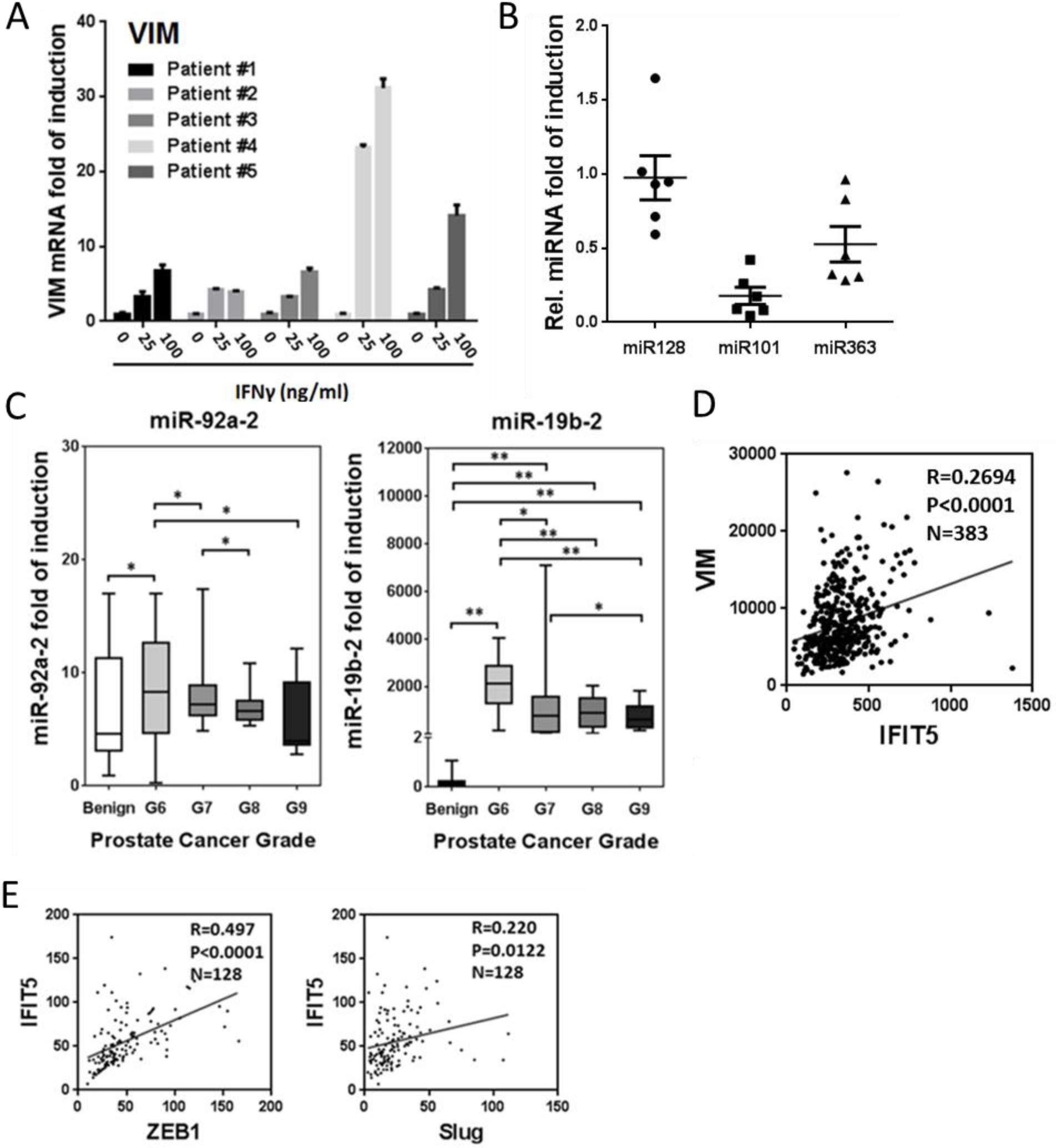
(A) Induction level of Vimentin mRNA expression in ex vivo culture of human PCa specimens (N=12) treated with IFNγ (0, 25 and 100 ng/ml) for 48hrs. (B) Induced relative fold change of miR-101, miR-128 and miR-363 expression in ex vivo culture of human PCa specimens treated with IFNγ (0, 25 and 100 ng/ml) for 48hrs. (C) Relative induction of miR-92a-2 and miR-19b-2 in human PCa specimens derived from different grades including benign (N=10), G6 (N=9), G7(N=9), G8(N=6) and G9(N=7) (*p<0.05, **p<0.0001). (D) Clinical correlation between IFIT5 and Vimentin (VIM) in PCa from TCGA PCa dataset. (E) Clinical correlation between IFIT5 and ZEB1 or Slug in renal cancer from TCGA renal cell carcinoma dataset.

### SUPPLEMENTAL TABLES

**Supplementary Table 1.** List of protein candidates derived from Mass Spectrometry after using precursor miR-363 to pull down protein complex in a variety of cell line pairs (LAPC4, RWPE1 and C4-2) with or without DAB2IP expression

| Cell line | DAB2IP-/DAB2IP+ Ratio | PSMs | Peptide Seqs | % Coverage | S. COUNTS |  | S. INDEX (MIC SIn) |  |
| --- | --- | --- | --- | --- | --- | --- | --- | --- |
|  |  |  |  |  | DAB2IP+ | DAB2IP- | DAB2IP+ | DAB2IP- |
| IFIT5_HUMAN Interferon-induced protein with tetratricopeptide repeats 5 OS=Homo sapiens GN=IFIT5 PE=1 SV=1 |  |  |  |  |  |  |  |  |
| LAPC4 | LAPC4-KD Only | 20 | 12 | 22.40 |  | 20.00 |  | 1.57E-06 |
| RWPE1 | 1.28 | 152 | 18 | 39.60 | 62.50 | 90.50 | 2.63E-05 | 3.37E-05 |
| C4-2 | C4-2Neo Only | 3 | 2 | 3.70 |  | 2.99 |  | 3.82E-07 |
| B9A067_HUMAN Mitochondrial inner membrane protein OS=Homo sapiens GN=IMMT PE=2 SV=2 |  |  |  |  |  |  |  |  |
| LAPC4 | 0.11 | 5 | 5 | 8.60 | 4.00 | 1.00 | 1.22E-07 | 1.39E-08 |
| RWPE1 | 0.28 | 5 | 3 | 7.30 | 3.00 | 2.00 | 3.57E-07 | 9.96E-08 |
| C4-2 | 0.17 | 58 | 31 | 36.80 | 49.05 | 7.85 | 6.29E-06 | 1.05E-06 |
| SNUT1_HUMAN U4/U6.U5 tri-snRNP-associated protein 1 OS=Homo sapiens GN=SART1 PE=1 SV=1 |  |  |  |  |  |  |  |  |
| LAPC4 | 0.40 | 14 | 10 | 14.80 | 9.00 | 5.00 | 2.37E-07 | 9.43E-08 |
| RWPE1 | 0.90 | 17 | 10 | 13.50 | 8.00 | 9.00 | 7.93E-07 | 7.11E-07 |
| C4-2 | 0.31 | 12 | 7 | 11.90 | 9.00 | 3.00 | 5.98E-07 | 1.88E-07 |
| ADT2_HUMAN ADP/ATP translocase 2 OS=Homo sapiens GN=SLC25A5 PE=1 SV=7 |  |  |  |  |  |  |  |  |
| LAPC4 | 0.32 | 24 | 9 | 28.90 | 12.96 | 10.97 | 5.78E-06 | 1.82E-06 |
| RWPE1 | 0.38 | 13 | 8 | 28.20 | 6.50 | 5.50 | 6.89E-06 | 2.63E-06 |
| C4-2 | 0.35 | 18 | 11 | 28.90 | 11.02 | 6.00 | 8.09E-06 | 2.87E-06 |
| PARP1_HUMAN Poly [ADP-ribose] polymerase 1 OS=Homo sapiens GN=PARP1 PE=1 SV=4 |  |  |  |  |  |  |  |  |
| LAPC4 | 0.68 | 135 | 40 | 40.80 | 73.00 | 62.00 | 3.38E-06 | 2.31E-06 |
| RWPE1 | 0.48 | 10 | 5 | 6.90 | 6.00 | 4.00 | 3.33E-07 | 1.61E-07 |
| C4-2 | 0.62 | 22 | 22 | 23.10 | 16.00 | 6.00 | 1.04E-06 | 6.42E-07 |
| XRCC5_HUMAN X-ray repair cross-complementing protein 5 OS=Homo sapiens GN=XRCC5 PE=1 SV=3 |  |  |  |  |  |  |  |  |
| LAPC4 | 0.36 | 29 | 16 | 22.80 | 18.98 | 9.99 | 6.73E-07 | 2.45E-07 |
| RWPE1 | 0.17 | 35 | 17 | 24.60 | 26.99 | 9.00 | 3.54E-06 | 6.10E-07 |
| C4-2 | C4-2D2 Only | 6 | 6 | 8.30 | 6.00 |  | 4.04E-07 |  |
| SFPQ_HUMAN Splicing factor, proline- and glutamine-rich OS=Homo sapiens GN=SFPQ PE=1 SV=2 |  |  |  |  |  |  |  |  |
| LAPC4 | 0.54 | 176 | 34 | 46.10 | 115.41 | 59.73 | 1.50E-05 | 8.10E-06 |
| RWPE1 | 0.69 | 11 | 6 | 12.00 | 5.00 | 6.00 | 7.40E-07 | 5.14E-07 |
| C4-2 | 0.58 | 27 | 14 | 19.80 | 15.96 | 10.99 | 2.57E-06 | 1.49E-06 |
| ATPA_HUMAN ATP synthase subunit alpha, mitochondrial OS=Homo sapiens GN=ATP5A1 PE=1 SV=1 |  |  |  |  |  |  |  |  |
| LAPC4 | 0.32 | 20 | 12 | 22.80 | 13.76 | 5.88 | 8.06E-07 | 2.55E-07 |
| RWPE1 | 0.94 | 28 | 12 | 29.30 | 13.78 | 13.79 | 1.75E-06 | 1.65E-06 |
| C4-2 | 0.88 | 97 | 25 | 41.60 | 49.17 | 46.19 | 1.13E-05 | 1.00E-05 |
| VDAC2_HUMAN Voltage-dependent anion-selective channel protein 2 OS=Homo sapiens GN=VDAC2 PE=1 SV=2 |  |  |  |  |  |  |  |  |
| LAPC4 | 0.05 | 12 | 8 | 32.50 | 10.87 | 0.99 | 1.36E-06 | 6.52E-08 |
| RWPE1 | 0.52 | 7 | 4 | 15.20 | 3.96 | 2.97 | 2.34E-06 | 1.21E-06 |
| C4-2 | 0.13 | 36 | 11 | 48.80 | 27.70 | 7.92 | 2.40E-05 | 3.20E-06 |
| H4_HUMAN Histone H4 OS=Homo sapiens GN=HIST1H4A PE=1 SV=2 |  |  |  |  |  |  |  |  |
| LAPC4 | LAPC4-Con Only | 5 | 3 | 31.10 | 5.00 |  | 2.46E-06 |  |
| RWPE1 | 0.70 | 7 | 4 | 40.80 | 5.00 | 2.00 | 3.79E-06 | 2.64E-06 |
| C4-2 | 0.08 | 7 | 3 | 31.10 | 6.00 | 1.00 | 1.14E-05 | 9.59E-07 |
| CKAP4_HUMAN Cytoskeleton-associated protein 4 OS=Homo sapiens GN=CKAP4 PE=1 SV=2 |  |  |  |  |  |  |  |  |
| LAPC4 | 0.41 | 6 | 3 | 6.00 | 3.00 | 3.00 | 8.02E-08 | 3.28E-08 |
| RWPE1 | 0.73 | 28 | 13 | 27.20 | 16.00 | 12.00 | 1.93E-06 | 1.41E-06 |
| C4-2 | 0.33 | 23 | 14 | 28.10 | 15.97 | 6.97 | 1.77E-06 | 5.76E-07 |
| ILF2_HUMAN Interleukin enhancer-binding factor 2 OS=Homo sapiens GN=ILF2 PE=1 SV=2 |  |  |  |  |  |  |  |  |
| LAPC4 | 1.25 | 303 | 21 | 48.20 | 140.00 | 163.00 | 1.31E-04 | 1.63E-04 |
| RWPE1 | 1.09 | 142 | 20 | 57.70 | 70.00 | 73.00 | 5.76E-05 | 6.29E-05 |
| C4-2 | 1.47 | 28 | 8 | 20.00 | 13.00 | 15.00 | 2.89E-06 | 4.27E-06 |
| ACACA_HUMAN Acetyl-CoA carboxylase 1 OS=Homo sapiens GN=ACACA PE=1 SV=2 |  |  |  |  |  |  |  |  |
| LAPC4 | 1.19 | 47 | 28 | 14.40 | 25.88 | 20.92 | 1.95E-07 | 2.32E-07 |
| RWPE1 | 1.31 | 820 | 115 | 58.50 | 174.15 | 259.86 | 2.10E-05 | 2.74E-05 |
| C4-2 | 8.98 | 16 | 122 | 43.40 | 5.94 | 9.44 | 8.89E-08 | 7.98E-07 |
| I3L1L3_HUMAN Myb-binding protein 1A (Fragment) OS=Homo sapiens GN=MYBBP1A PE=4 SV=1 |  |  |  |  |  |  |  |  |
| LAPC4 | LAPC4-Con Only | 22 | 19 | 17.90 | 22.00 |  | 6.31E-07 |  |
| RWPE1 | 0.29 | 3 | 2 | 1.80 | 2.00 | 1.00 | 5.15E-08 | 1.47E-08 |
| C4-2 | C4-2D2 Only | 2 | 2 | 1.60 | 2.00 |  | 3.41E-08 |  |

**Supplemental Table 2.**
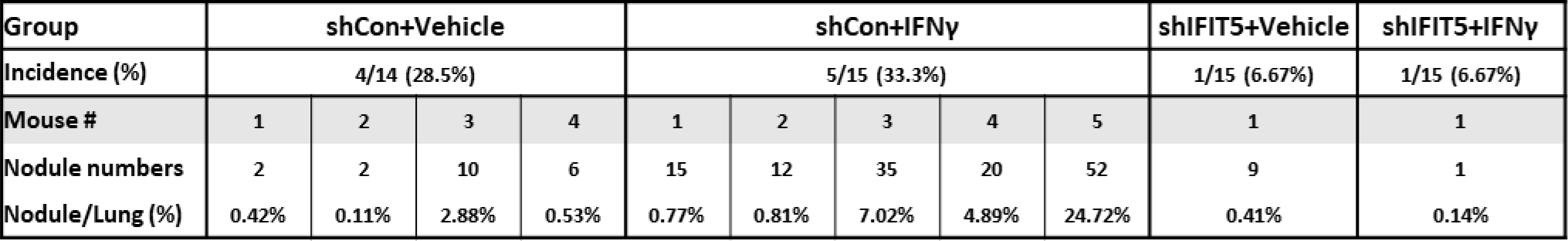
The lung metastatic tumor incidence and nodule formation in SCID mice receiving intravenous injection of PC3 cells through tail vein.

| Group | shCon+Vehicle | | | | shCon+IFNγ | | | | | shIFIT5+Vehicle | shIFIT5+IFNγ |
| --- | --- | --- | --- | --- | --- | --- | --- | --- | --- | --- | --- |
| Incidence (%) | 4/14 (28.5%) |  |  |  | 5/15 (33.3%) |  |  |  |  | 1/15 (6.67%) | 1/15 (6.67%) |
| Mouse # | 1 | 2 | 3 | 4 | 1 | 2 | 3 | 4 | 5 | 1 | 1 |
| Nodule numbers | 2 | 2 | 10 | 6 | 15 | 12 | 35 | 20 | 52 | 9 | 1 |
| Nodule/Lung (%) | 0.42% | 0.11% | 2.88% | 0.53% | 0.77% | 0.81% | 7.02% | 4.89% | 24.72% | 0.41% | 0.14% |

**Supplemental Table 3.**
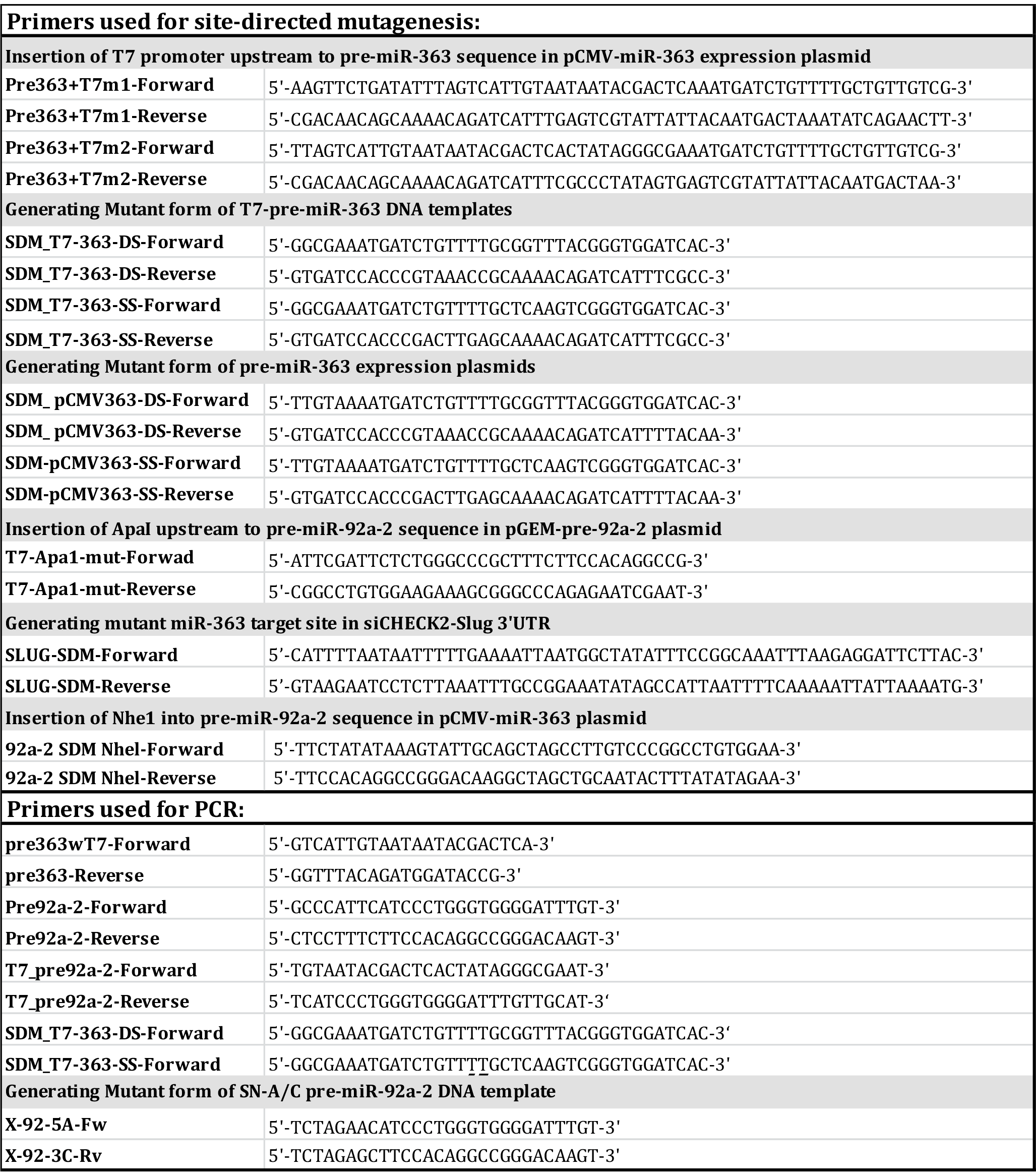
Primers designed for site-directed mutagenesis and PCR.

### SUPPLEMENTAL INFORMATION

#### Plasmid constructions

##### Luciferase reporter plasmid siCHECK2-Slug3’UTR and psiCHECK2-Slug3’UTR-Mut^363^

psiCHECK2-dual luciferase reporter construct (Promega) containing the wild type 3’UTR sequence of human SNAI2/Slug (psiCHECK2-Slug3’UTR-WT) was obtained from Dr. Jin-Tang Dong (Emory University). The Slug 3’UTR contains one putative target site of miR-363. We generated a psiCHECK2-Slug3’UTR-Mut^363^ plasmid by mutating the miR-363 target site using site-directed mutagenesis kit.

##### pCMV-miR-363 expression vector

The pCMV-miR-363 expression plasmid initially purchased from Origene contains both pre-miR-92a-2 and pre-miR-363 sequence. In order to isolate pre-miR-363 sequence, we generated a Nhel cut site in the 5’-end of pre-miR-92a-2 using site-directed mutagenesis and excised pre-miR-363 sequence by using both NheI and MluI enzyme then inserted into the pCMV-miRNA empty vector to generate miR-363 expression plasmid (Native pre-miR-363).

##### SSMut and DSMut precursor miRNA-expression plasmid constructs

We used Native miR-363 expressing plasmid (Origene) as a template to generate mutant pre-miR-363 with 5’-six nucleotides single stranded overhang (SSMut pre-miR-363) or double-stranded blunt end (DSMut pre-miR-363) constructs using site-directed mutagenesis kit. We used Native miR-101 and miR-128 expressing plasmid (Genecopoeia) as templates to generate mutant pre-miR-101 with 5’-nine nucleotides single stranded overhang (SS9Mut pre-miR-101) or double-stranded blunt end (DSMut pre-miR-101) constructs, as well as mutant pre-miR-128 with 5’-six nucleotides single stranded overhang (SS6Mut pre-miR-128) or double-stranded blunt end (DSMut pre-miR-128) constructs using site-directed mutagenesis.

##### Plasmid construct for *in vitro* transcription of Native and Mutant pre-miR-363 RNA molecules

A sequence of T7 promoter was inserted into the upstream of Native pre-miR-363 plasmid using two-step site-directed mutagenesis. This plasmid was further used as a template to generate T7-SS6Mut pre-miR-363 and T7-DSMut pre-miR-363 constructs using site-directed mutagenesis kit. Subsequently, these DNA templates were PCR amplified and cloned into pGEM-T_Easy_ vector (Promega) for sequencing. After DNA sequencing confirmation, they were subjected to *in vitro* transcription to generate Native, SSMut and DSMut pre-miR-363 RNA molecules.

##### Plasmid construct for *in vitro* transcription of pre-miR-92a-2 RNA molecules

The pre-miR-92a-2 sequence was PCR amplified and cloned into the downstream of T7 promoter in the pGEM-T_Easy_ vector. In order to generate a template for *in vitro* transcription, one ApaI site in addition to the internal ApaI (5 nucleotides downstream from the T7 promoter) was created from 4 nucleotides upstream from the pre-miR-92a-2 sequence. After ApaI cleavage and gel purification, the plasmid was re-ligated to generate T7-pre-miR-92a-2 DNA template that is used for *in vitro* transcription for produce pre-miR-92a-2 RNA molecules.

##### Purification of recombinant IFIT5 protein (rIFIT5)

The pET-28a-6XHis-3XFlag-IFIT5 plasmid given by Dr. Collins was transformed and expressed in BL21 competent cells. BL21 cells were induced with 0.2 mM isopropyl β-D-thiogalactopyranoside (IPTG) and grown overnight at 22°C. After spin down at 10,000 rpm for 10min at 4°C, cell pellet was re-suspended in Buffer A containing 20 mM HEPES, pH 7.5, 150 mM NaCl, 5 mM imidazole, 1 mM tris(2-carboxyethyl) phosphine (TCEP), 10% glycerol, with protease inhibitor cocktail (Thermo Scientific). Suspended BL21 cells were lysed by sonication on ice for 1 hr with 5-sec interval. After centrifugation at 35,000xg for 1 hr at 4°C, the cleared lysate was applied to a Ni-charged HiTrap FF column (GE Healthcare), and bound His-tagged IFIT5 proteins were denatured by washing with Buffer A containing 6 M guanidine hydrochloride to remove bound RNA. Afterwards, the RNA-free IFIT5 protein was refolded on the column by washing with Buffer A. Refolded IFIT5 was eluted in Buffer A containing 500 mM imidazole and final elute was dialyzed using 10K slide-A-Lyzer dialysis cassette (Thermo Scientific). The purity of recombinant IFIT5 protein was confirmed by western blot analysis for subjected to *in vitro* RNA degradation assay.

